# RNA Pol II Length and Disorder Enable Cooperative Scaling of Transcriptional Bursting

**DOI:** 10.1101/825299

**Authors:** Porfirio Quintero-Cadena, Tineke L. Lenstra, Paul W. Sternberg

**Affiliations:** California Institute of Technology, Division of Biology and Biological Engineering, Pasadena, CA; The Netherlands Cancer Institute, Oncode Institute, Division of Gene Regulation, Amsterdam, the Netherlands

**Keywords:** Transcriptional bursting, RNA Pol II, CTD length, Phase separation

## Abstract

RNA Polymerase II contains a disordered C-terminal domain (CTD) whose length enigmatically correlates with genome size. The CTD is crucial to eukaryotic transcription, yet the functional and evolutionary relevance of this variation remains unclear. Here, we use smFISH, live imaging, and RNA-seq to investigate how CTD length and disorder influence transcription. We find that length modulates the size and frequency of transcriptional bursting. Disorder is highly conserved and mediates CTD-CTD interactions, an ability we show is separable from protein sequence and necessary for efficient transcription. We build a data-driven quantitative model, simulations of which recapitulate experiments and support CTD length promotes initial polymerase recruitment to the promoter but slows down its release from it, and that CTD-CTD interactions enable promoter recruitment of multiple polymerases. Our results reveal how these tunable parameters provide access to a range of transcriptional activity, offering a new perspective for the mechanistic significance of CTD length and disorder in transcription across eukaryotes.

## 1. Introduction

In eukaryotes, the RNA Polymerase II complex that transcribes protein-coding genes is typically composed of 12 subunits (Hantsche and Cramer, 2017). The largest and catalytic subunit RPB1 contains a repetitive and unstructured C-terminal Domain (CTD) that is a major factor for establishing critical protein-protein interactions throughout transcription and downstream processes (Harlen and Churchman, 2017).

Each of the heptad amino acid repeats in the CTD, whose number ranges from 5 in *Plasmodium yoelii* to 60 in *Hydra* (Chapman et al., 2008; Yang and Stiller, 2014), can be subject to post-translational modifications that regulate its physical interactions and consequently RNA Pol II function (Eick and Geyer, 2013; Harlen and Churchman, 2017). The CTD’s repetitive nature most likely arose in the last eukaryotic common ancestor (Yang and Stiller, 2014), and its length appears to enigmatically correlate with genome size (Chapman et al., 2008; Yang and Stiller, 2014). What is the role of CTD length variation in transcription?

Truncating the number of CTD repeats impacts cell growth and animal development, with a minimal number required for viability (Nonet et al., 1987; Bartolomei et al., 1988; West and Corden, 1995; Gibbs et al., 2017; Lu et al., 2019), and reduces the transcriptional output from enhancer responsive genes (Allison and Ingles, 1989; Scafe, C; Young, 1990; Gerber et al., 1995; Aristizabal et al., 2013). Enhancers physically interact with promoters via protein-protein interactions to activate transcription in bursts of activity (Chubb et al., 2006; Bartman et al., 2016; Chen et al., 2018). Given that mRNA output decays rapidly with increasing separation between enhancers and promoters (Dobi and Winston, 2007; Quintero-Cadena and Sternberg, 2016), an intriguing possibility is that CTD expansion facilitates enhancer function over physical distances to promoters that scale with genome size (Allen and Taatjes, 2015).

Increasingly relevant for the understanding of biological phenomena, liquid-liquid phase separation is an emerging signature of proteins with disordered, repetitive domains (Banani et al., 2017; Shin and Brangwynne, 2017). Low complexity domains that exhibit this property appear to be abundant in nuclear proteins, including the CTD and major transcription factors (Cho et al., 2018; Chong et al., 2018; Qiu et al., 2018; Shin et al., 2018). CTD length has also been implicated in its ability to form (Boehning et al., 2018) and bind phase-separated droplets, with a minimum threshold that parallels its viability requirement (Kwon et al., 2013). In addition, the interaction of the CTD with these droplets can be dynamically modulated by phosphorylation (Kwon et al., 2013; Chong et al., 2018; Boehning et al., 2018; Cho et al., 2018; Nair et al., 2019), a major post-translational modification that precedes transcription initiation (Payne et al., 1989; Svejstrup et al., 1997). In light of these observations, phase separation provides an appealing framework to explain certain transcriptional phenomena. From this perspective, the CTD could provide a bridge for the RNA polymerase to dynamically participate in multi-molecular assemblies of transcription factors and DNA loci that facilitate the function of highly active enhancers (Hnisz et al., 2017).

Extensive investigations have revealed many roles of CTD sequence and post-translational modifications (Eick and Geyer, 2013; Harlen and Churchman, 2017). On the other hand, the functional and evolutionary relevance of CTD length and the mechanism by which it influences transcription have not been systematically investigated.

Here, by quantitatively analyzing snapshots and dynamics of transcription in budding yeast, we show that CTD length can modulate transcription burst size and frequency. We strengthen the evolutionary relevance of the CTD’s long disorder, and provide evidence that its role in transcription is separable from amino acid sequence. Specifically, we demonstrate that the function of the CTD’s long disorder can be supplemented by similarly unstructured protein domains. These proteins can interact with and recruit others of their kind, an ability that is necessary for efficient transcription *in vivo*. We use these features, together with known CTD protein-protein interactions, to construct an integrative and quantitative model that explains how CTD length influences the dynamics of eukaryotic transcription.

## 2. Results

### 2.1. CTD is enriched in disordered amino acids and its length inversely correlates with gene density across eukaryotes

The CTD of representative species has been shown to be a random coil (Portz et al., 2017), suggesting this structural feature is relevant for its function and could itself be used to identify it. We sought to learn whether this was a common signature in all known protein sequences. We searched for RPB1 protein sequence homologs, the CTD-bearing catalytic subunit of RNA Pol II (Figure 1A, top). We recovered 542 unique sequences from 539 species and 338 genera, whose length varies from 1374 to 3055 amino acids (Figure 1B), with some evident bias that likely stems from the limited availability of genome sequences.

**Figure 1:**
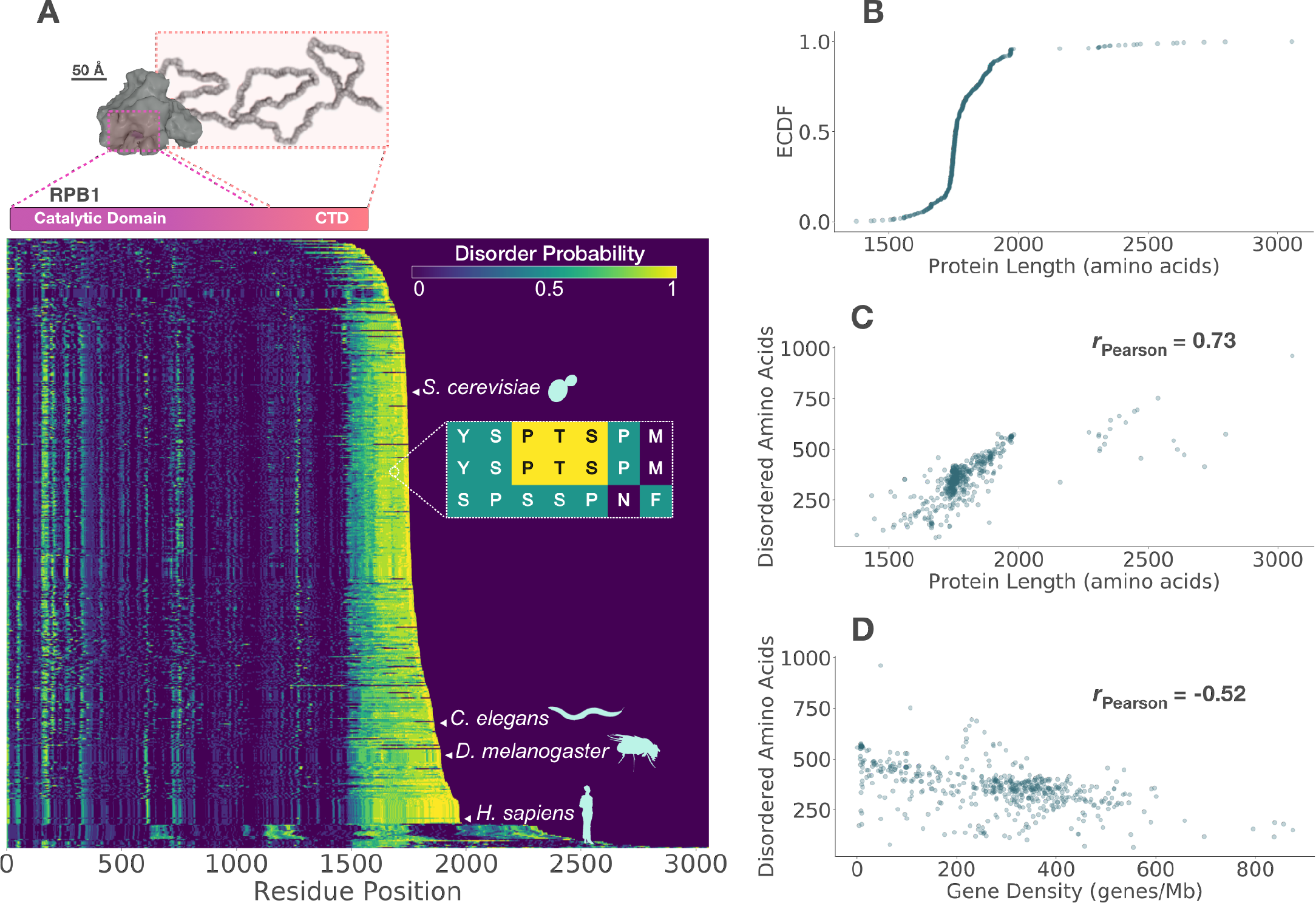
RPB1, the catalytic subunit of RNA Polymerase II, contains an unstructured C-terminal Domain (CTD) whose length correlates with gene density across eukaryotes. (A) Top: Cartoon of 12 subunit RNA Pol II complex with an unstructured CTD drawn to scale, adapted from 1Y1W.PBD (Kettenberger et al., 2004; Portz et al., 2017). RPB1 is highlighted in pink and its CTD in orange. Bottom: distribution of disorder probability along RPB1 sequence homologs sorted by length. Representative species are highlighted. Inset illustrates residue coloring by disorder probability. (B) Empirical cumulative distribution function (ECDF) of RPB1 lengths. (C) RPB1 length positively correlates with number of predicted disordered amino acids and inversely with gene density (D). Pearson correlation coefficient is shown for each pair of variables.

We computed the disorder probability per amino acid along each protein sequence. We found that with few exceptions, the C-termini of RPB1 sequences is enriched in disordered amino acids (Figure 1A, bottom). Protein length is positively correlated with the number of disordered amino acids (Figure 1C). As noted in previous reports (Chapman et al., 2008; Yang and Stiller, 2014), these C-terminal sequences are enriched in amino acids from the heptad repeat YSPTSPS (Figure S1A). Like amino acid content, overall charge, aromaticity and hydrophobicity have a compact distribution (Figure S1B-D).

CTD length has been shown to correlate with genome size for a few representative species that span a wide range of genome sizes (Chapman et al., 2008; Yang and Stiller, 2014), from 1×10^7^ bp in yeast to 3×10^9^ bp in human. To systematically investigate the generality of this phenomenon, we compiled a list of genome sizes and their estimated gene number. We then computed the gene density (genes per megabase of DNA) for each species, in order to account for long stretches of non-coding DNA that are more common in large genomes. The number of disordered amino acids in RPB1 homologs inversely correlated with gene density (Figure 1D): sparse genomes tend to have polymerases with longer CTDs.

This systematic characterization of RPB1 sequences builds on previous extensive reports (Chapman et al., 2008; Yang and Stiller, 2014) and highlights three important features of the CTD. First, protein disorder is a highly conserved and likely functionally relevant feature of the CTD. Second, RPB1 length variation mostly originates from the number of disordered amino acids in its C-terminus. Third, CTD length is inversely correlated with gene density.

### 2.2. CTD length modulates transcription burst size and frequency

We sought to understand the role of CTD length using *Saccharomyces cerevisiae* as our model. Wild-type yeast CTD contains 26 heptad repeats (CTDr). We generated strains in which the genomic copy of RPB1 was engineered to have 14, 12, 10, 9, and 8 CTDr, the minimum required for viability in yeast (West and Corden, 1995). Consistent with previous reports (Nonet et al., 1987; West and Corden, 1995), the growth rate of these strains was compromised by CTD truncation. The magnitude of the decrease in growth rate increased with decreasing CTD length (Figure 2A); while 14 and 12 CTDr strains have only a subtle growth phenotype, the decrease in growth rate becomes evident with 10 CTDr and progressively larger with 9 and 8 CTDr (Figure 2A, inset).

**Figure 2:**
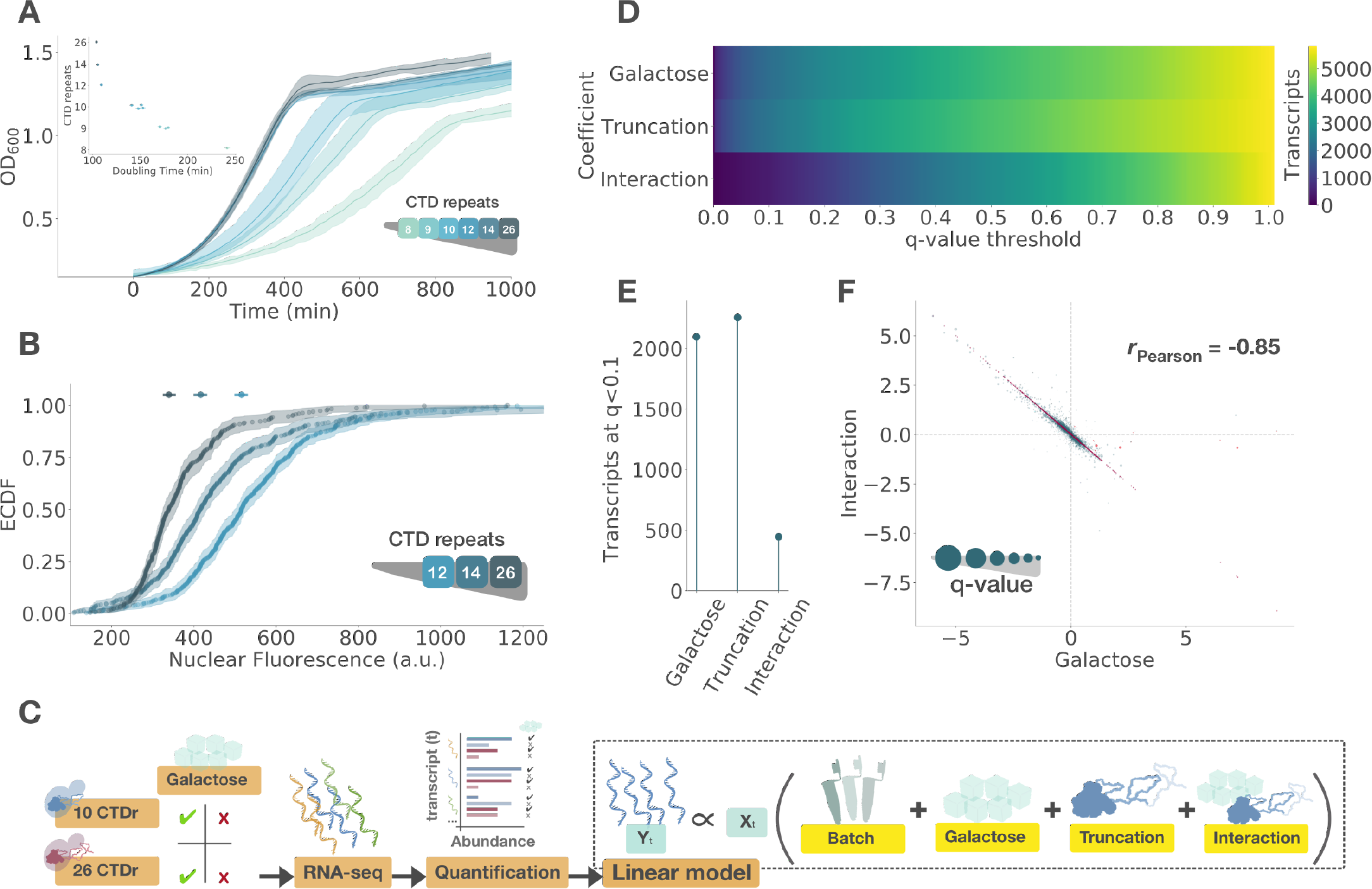
CTD truncation reduces growth rate, increases nuclear Pol II concentration and antagonizes transcription activation genome-wide. (A) Mean optical density (OD) over time of *S. cerevisiae* strains with 26 (wild-type), 14, 12, 10, 9, and 8 CTD repeats (CTDr). Shaded area shows the range of measured ODs at each time point from three biological replicates per line. Inset shows mean doubling times (DT) with standard error by line. (B) Empirical cumulative distribution function (ECDF) of mScarlet-RPB1 nuclear fluorescence in strains with 26, 14, and 12 CTDr from three biological replicates. Shaded area is bootstrapped 99% confidence interval (CI) and top markers show median with 99% CI. (C) Experimental design to measure the transcriptomic phenotype of CTD truncation and its influence on the transcriptional response after 2 hours of galactose induction via RNA-seq from three biological replicates. A linear model was used to fit the RNA-seq data, where each coefficient estimates the influence on each measured transcript of batch effects, galactose induction, CTD truncation and its interaction with induction. For each coefficient, (D) cumulative distribution, in number of transcripts, of false discovery rates (q-values) and (E) number of differentially expressed transcripts detected at a q-value threshold of 0.1. (F) Comparison of the log fold-change of each transcript resulting from galactose induction and its interaction with CTD truncation. Red points show the positions on the diagonal *x* = *y*. Marker size of each point is inversely proportional to the q-value of the interaction (*ms* = −*log*(*q*_*int*_)); dotted lines reference no change at zero and the Pearson correlation is indicated. Direct targets of GAL4 listed in Lesurf et al. (2016) are plotted in orange.

RNA Pol II is mostly present in the nucleus, imported as a fully assembled complex (Boulon et al., 2010; Czeko et al., 2011). We hypothesized that nuclear polymerase becoming rate-limiting could explain the observed phenotypes, given that the CTD has been linked to nuclear import (Carre and Shiekhattar, 2011). We fused the fluorescent protein mScarlet to RPB1, and unexpectedly found that CTD truncation increased its nuclear levels (Figure 2B). This effect is presumably explained by the requirement of the CTD for ubiquitination (Huibregtse et al., 1997; Somesh et al., 2005) and the resulting accumulation of the protein complex. We were unable to fuse mScarlet to shorter CTD strains; however, this trend suggested the polymerase does not become rate-limiting upon CTD truncation.

We then asked how CTD length influences transcription. We focused on the galactose transcriptional response, because like other inducible pathways it has been shown to be sensitive to CTD truncations (Allison and Ingles, 1989; Scafe, C; Young, 1990) and involves the differential expression of over 2000 transcripts (Figure 2E), about a third of the yeast’s transcriptome. We measured the transcriptional phenotype of CTD truncation using RNA-seq, comparing wild-type with the 10CTDr strain with and without galactose (Figure 2C). We fitted these data to a linear model that allowed us to estimate the individual contributions, for each measured transcript, of four components in the experiment: batch effects, galactose induction, CTD truncation, and its interaction with galactose induction (Figure 2C, right).

We identified over 2000 transcripts affected by CTD truncation at a q-value threshold of 0.1, from a distribution of q-values that indicates a strong transcriptional phenotype (Figure 2D,E). A significant proportion of the galactose responding transcripts exhibited a statistical interaction with CTD truncation. This effect revealed a surprisingly specific, globally antagonistic relationship between two components: 1) galactose induction and 2) the interaction of truncation with galactose induction (Figure 2F). In other words, CTD truncation reduced the magnitude of change in abundance of most transcripts upon galactose induction.

We sought to understand the source of this antagonism by visualizing transcription dynamics in living cells. We introduced 14 copies of the bacteriophage sequence PP7 in the 5’ UTR of GAL10, a strongly galactose responsive gene. Each of the PP7 repeats forms an RNA hairpin that can be bound by a pair of PP7 coat proteins fused to GFP (Coulon et al. (2014); Lenstra et al. (2015), Figure 3A, top). This system allowed us to visualize the dynamics of GAL10 transcription upon galactose induction as fluorescence bursts arising from the transcription site (TS; Figure 3A, bottom). We found that expressing TS intensity as the ratio of spot to mean nuclear fluorescence could reliably account for the differences in PP7-GFP levels observed between strains (Figure S2D-H).

**Figure 3:**
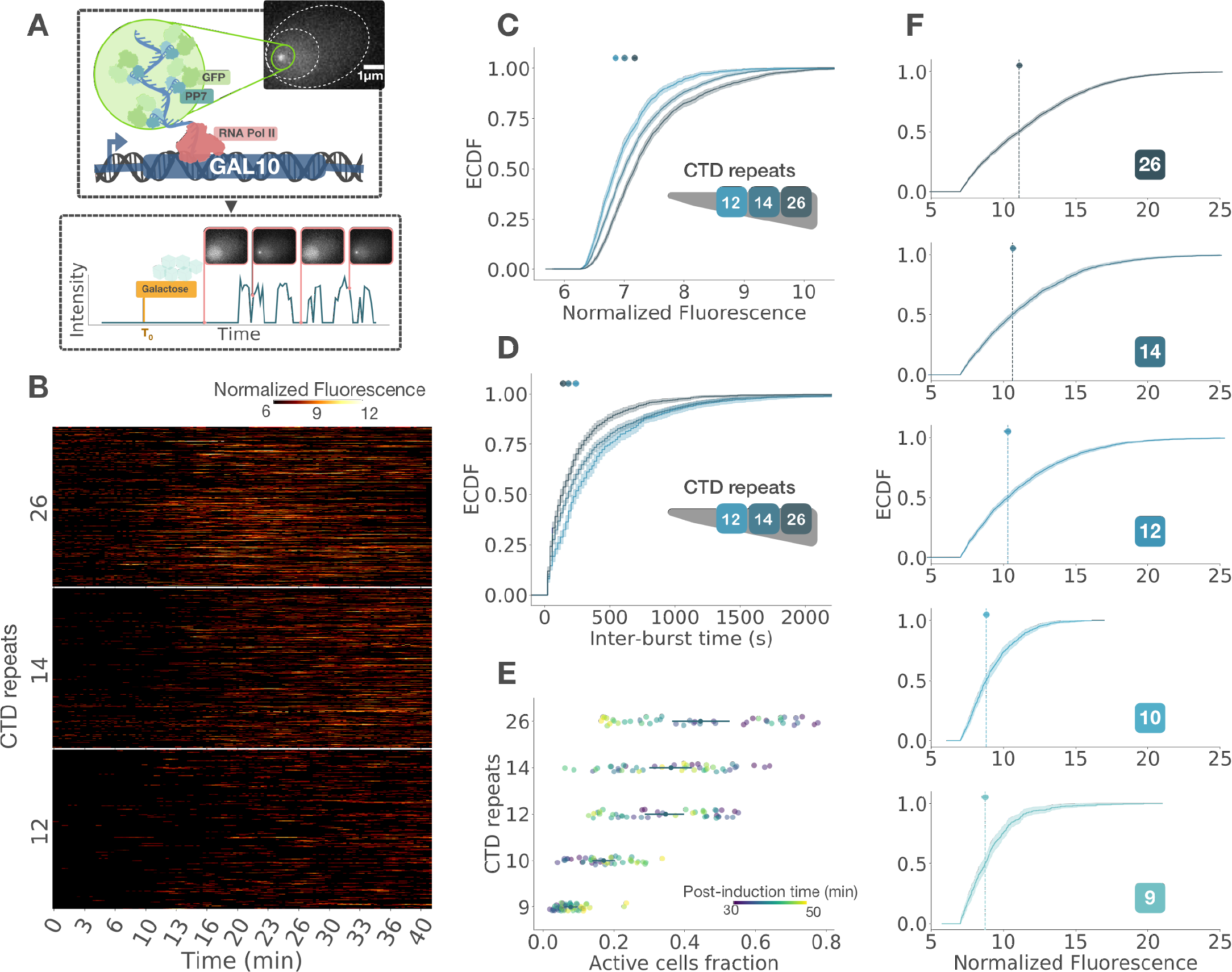
CTD length modulates transcription burst size and frequency. (A) Experimental strategy used to observe transcription dynamics in live cells. The 5’ end of a single allele of galactose-inducible GAL10 is tagged with RNA hairpins that are bound by nuclear expressed GFP-PP7. This protein fusion results in a fluorescent spot in the cell nucleus upon transcription activation, whose fluorescence through time is recorded to investigate transcription dynamics. (B) Transcriptional traces by cell of strains with 26 (wild-type), 14, and 12 CTD repeats (CTDr) from three biological replicates. These are related to burst size and frequency through (C) the empirical cumulative distribution function (ECDF) of transcription site fluorescence intensities and the ECDF of inter-burst times, in seconds (D). Shaded area is bootstrapped 99% confidence interval (CI) and top markers show median with 99% CI. (E) Fraction of active cells per field of view from two biological replicates measured from high-laser-power snapshots of strains with 26, 14, 12, 10, and 9 CTDr. Middle points indicate mean with bootstrapped 99% CI. Color indicates time after galactose induction. (F) ECDFs of normalized fluorescence intensities of transcription bursts from these snapshots. Vertical dotted lines indicate median and shaded area bootstrapped 99% CI. Number of CTD repeats is indicated in the lower right corner of each plot.

Given the burstiness of transcription, we hypothesized that two parameters could play a role in the diminished transcriptional output, namely burst size and frequency. We measured transcription fluorescence traces for 26 (wild-type), 14, and 12 CTDr strains (Figure 3B). CTD truncation decreased the intensity of fluorescence bursts (Figure 3C), suggesting a decrease in burst s ize. Also, truncation increased the time interval between bursts (Figure 3D), which is closely related to burst frequency. We could similarly observe this frequency decay by looking at whether the average cell was active or inactive over time for each strain (Figure S3A). These average traces also show that burst frequency remained roughly constant after activation, only declining towards the end likely due to photobleaching and mRNA-bound PP7-GFP nuclear export. The autocorrelation of normalized intensity traces increased in amplitude with CTD truncation, similarly supporting a decrease in burst frequency (Figure S3B-D). Differences in burst duration measured from this analysis were more subtle. This potential ambiguity could mean that burst duration is independent of size, or that the observed decay in TS intensity was influenced by the decay in burst frequency. Overall, the live transcription measurements suggested CTD length can simultaneously modulate burst size and frequency.

The transcriptional activity in strains with shorter CTDs was too weak to be visualized in these movies without incurring in phototoxic illumination. To circumvent this problem, we took a single snapshot per field of view with maxium laser intensity during a 20 minute window after 30 minutes of galactose induction. This approach additionally allowed us to obtain a better estimate of TS fluorescence. The fraction of active cells per field of view was impacted by CTD truncation (Figure 3E). We observed a transition similar to the growth phenotype in this assay, where the magnitude of the effect progressively increased with CTD truncation. We also observed a consistently moderate shift in the distributions of TS fluorescence with CTD truncation (Figure 3F). The comparatively small magnitude of this decrease suggested the decay in fraction of active cells is primarily driven by burst frequency. From these measurements, we conclude that CTD length can modulate both the size and the frequency of transcriptional bursting.

### 2.3. Fusion to disordered proteins can rescue the function of a CTD-truncated RNA Pol II

Given the conservation of protein disorder (Figure 1A) and recent evidence that the CTD can form and interact with phase separated droplets (Kwon et al., 2013; Chong et al., 2018; Boehning et al., 2018; Cho et al., 2018; Nair et al., 2019), we hypothesized that the function of the CTD’s long disorder could be supplied by other proteins of similar chemical and structural features. We tested this idea by fusing the low complexity domains (LCD) of the human proteins FUS and TAF15, which are not present in the yeast genome, to the C-terminus of a 10CTDr truncated RPB1. These LCDs contain neither a known nuclear localization sequence (Gal et al., 2011; Marko et al., 2012) nor ubiquitination sites (Mertins et al., 2013) that could supplement CTD function in a predictable manner; they are similar in amino acid composition, particularly in the frequency of tyrosines, but share little sequence similarity with the CTD (Figure S4A-C).

Astonishingly, strains carrying either protein fusion showed an improved growth rate over the 10CTDr strain. Fusion to FUS LCD progressively rescued growth rates of strains with 9 and 8 CTDr. Furthermore, strains with 7 and 6 CTDr remained viable when fused to FUS, surpassing the minimum requirement of 8 CTDr alone (Figure 4A). This supression was particularly striking because the CTD has been extensively mutated, typically with detrimental effects to transcription or downstream processes. This new minimal length is also noticeably close to the four heptad repeats that directly contact Mediator in an assembled preinitiation complex (Robinson et al., 2012, 2016).

**Figure 4:**
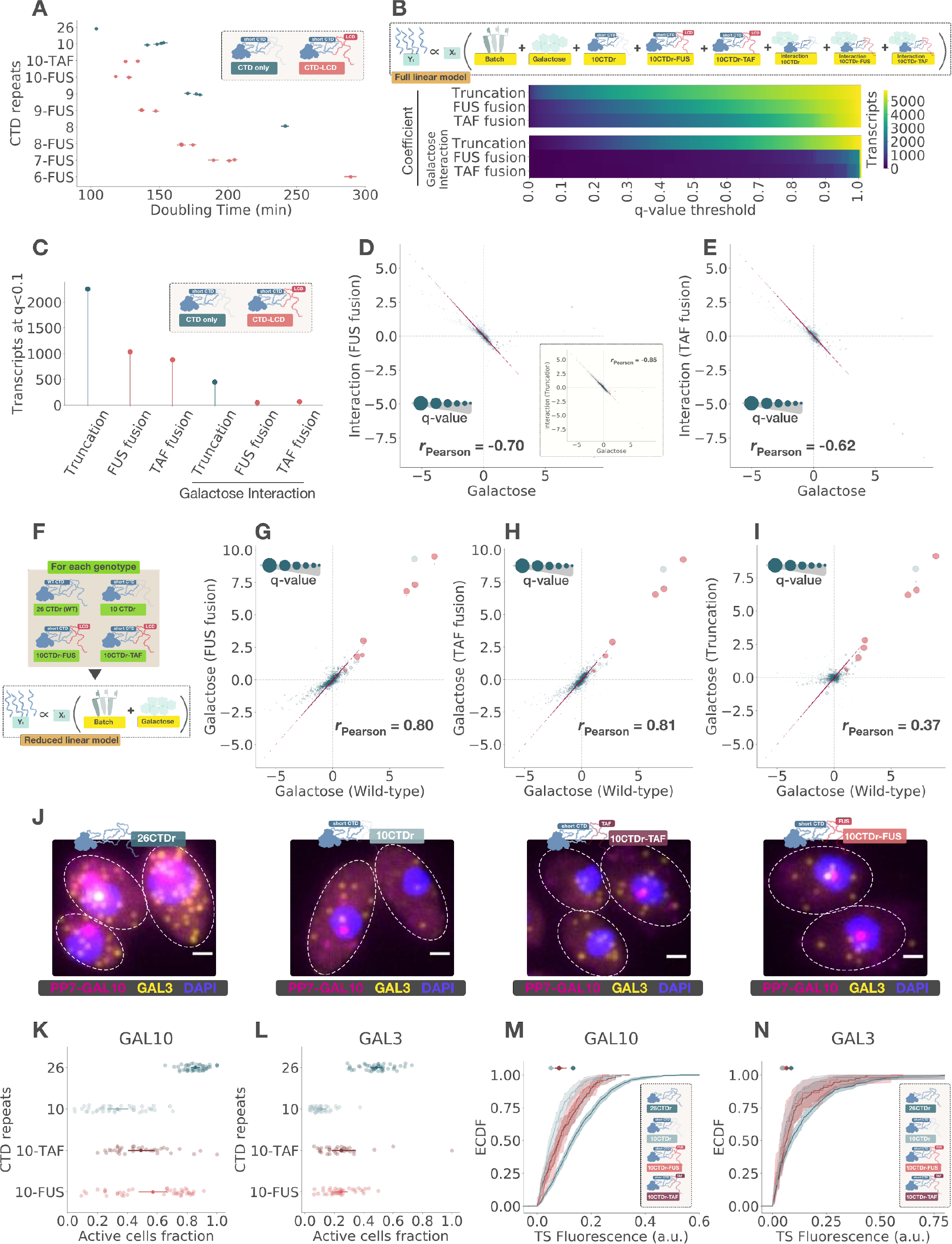
Fusion of the low-complexity domain (LCD) of FUS or TAF15 to a truncated polymerase can rescue its function and reduce the CTD length required to support cell growth. (A) Comparison of doubling times (DT) of strains with wild-type and decreasing number of CTD repeats (CTDr) with and without fusion to the LCD of FUS or TAF15. Individual points come from independent lines when available and indicate mean DTs with standard error estimated from three biological replicates per line. (B) Top: Linear model used to estimate the effect of truncation to 10CTDr, subsequent LCD fusion, and the interaction of each of these components with galactose induction. Bottom: Resulting cumulative distributions of q-values, in number of transcripts, for each coefficient, excluding galactose and batch effects from three biological replicates. (C) Number differentially expressed transcripts detected at a q-value threshold of 0.1. Comparison of the log fold-change of each transcript induced by galactose and its interaction with 10CTDr fused with FUS (D) and TAF15 (E) LCDs. Inset shows comparison with 10CTDr interaction alone. (F) Reduced linear model used to measure galactose induction in each strain individually. Comparisons of log fold-change of each transcript induced by galactose in wild-type and 10CTDr (G), 10CTDr-FUS (H), and 10CTDr-TAF15 (I). Red points show the positions on the diagonal *x* = *y*. Marker size of each point is inversely proportional to the q-value of the coefficient in the y-axis (*ms* = −*log*(*q*_*y*_)); dotted lines reference no change at zero and the Pearson correlation is indicated. Direct targets of GAL4 listed in Lesurf et al. (2016) are plotted in orange. (J) Representative images of smFISH with probes for a single allele of PP7-GAL10 and both alleles of GAL3 after two hours of galactose induction for 26CTDr (wild-type), 10CTDr, 10CTDr-TAF, and 10CTDr-FUS strains as indicated on top of each image. White dotted contours mark cell outlines. Scale bar is 1 *μ*m. Fraction of active cells per field of view for GAL10 (K) and GAL3 (L) measured from two biological replicates of smFISH. Mean with 99% bootstrapped confidence interval (CI) is shown on top of each group. Corresponding empirical cumulative distribution functions (ECDF) with 99% bootrsapped CI of transcription site intensities of GAL10 (M) and GAL3 (N). Medians with 99% CI are shown on top.

We next probed whether the improved growth phenotype originated from a transcriptional rescue. We used RNA-seq to compare the transcriptomes of the FUS and TAF15 rescued strains and their response to galactose induction with that of the wild-type and 10CTDr strains. Using principal component analysis, we observed LCD fusion results in transcriptomes in-between that of the wild-type and truncated strains, under induced and uninduced conditions (Figure S5A). As described in Figure 2B, we fitted the data to a linear model to identify the contributions to each transcript of galactose induction, CTD truncation, FUS or TAF15 LCD fusion to a 10CTDr truncated polymerase, and their interaction with galactose (Figure 4B, top). We found the number of differentially expressed transcripts resulting from CTD truncation decreased from 2256 to 1037 and 883 transcripts at a q-value threshold of 0.1 upon fusion to FUS or TAF15 LCDs, respectively (Figure 4C). More generally, the distribution of q-values resulting from CTD truncation shifted towards less significant values (Figure 4B, middle), a sign of considerable amelioration in the transcriptional phenotype. These LCD fusions additionally shifted the distribution of q-values of the interaction between CTD truncation and galactose induction (Figure 4B, bottom), abolishing the measurable effect at a q-value of less than 0.1 (Figure 4C). Moreover, the global antagonism of this interaction term with galactose induction nearly vanished (Figure 4D,E).

We interrogated these data further using other linear models, the simplest of which consists of independently measuring the galactose induction in each strain (Figure 4F). In this comparison, the galactose response of most transcripts in the rescued strains closely resembles that of the wild-type (Figure 4G,H), more than the 10CTDr alone (Figure 4I). The FUS and TAF15 transcriptional phenotypes, as measured using the full model (Figure 4B, top), were highly correlated (Figure S5B). Based on this observation, we fitted the data to a model in which we consider the two rescued strains as a single LCD group by pooling their transcriptomes together (Figure S5C). This model increased the number of transcripts that we can confidently call differentially expressed at a q-value threshold of 0.1 to 1392 (Figure S5D). This effect supports the transcriptional rescue occurs through a single pathway, and that there is a CTD sequence-dependent signature that remains shared among the three 10CTDr strains. This signature was evident from the high similarity between the truncation and LCD fusion coefficients (Figure S5E). From this experiment, we conclude that the sequence and long disorder of the CTD have separable roles in transcription, the latter of which can be supplemented by the similarly disordered LCDs of FUS or TAF15.

For unknown reasons, our live imaging system did not work with the FUS rescued strains. We circumvented this issue by using two-color Single-Molecule Fluorescence *in situ* Hybridization (smFISH). We used probes against the PP7 repeats, allowing us to detect mRNA from a single allele of GAL10, and against GAL3, which could detect RNA from both of its alleles (Figure 4J). The fraction of active cells consistently increased for both the single allele of GAL10 and the two alleles of GAL3 (Figure 4K,L). In addition, we observed an increase in TS fluorescence intensity of both rescued strains over the 10CTDr strain (Figure 4L,M), suggesting the fraction of active cells increased specifically because of an increased burst size.

Together, these results show the CTD’s long disorder can influence transcription in a way that depends on its chemical and structural properties rather than its precise amino acid sequence.

### 2.4. CTD, FUS and TAF15 LCDs can self-interact and this ability is necessary for efficient transcription

The CTD can bind the LCD of FUS and more strongly that of TAF15 (Kwon et al., 2013). TAF15 can also interact with components of the Mediator complex (Takahashi et al., 2011). However, the avidity of these interactions does not appear to correlate with the extent of rescue (Figure 4A). Other experiments have shown these LCDs are able to phase-separate (Kwon et al., 2013). These phases are thought to form as a result of intermolecular interactions that collectively drive droplet formation (Banani et al., 2017; Shin and Brangwynne, 2017). We hypothesized that FUS, TAF15, and the CTD could be involved in the recruitment of RNA Pol II to the TS via these self-interactions.

We first tested whether FUS and TAF15 variants, with tyrosine to serine misense mutations that make them significantly less able to bind phase-separated droplets *in vitro* (Kwon et al. (2013); data reproduced in Figure S4D), would fail to rescue the growth phenotype of a 10CTDr strain.

We observed a correlation between droplet binding rates and the extent of growth rescue upon fusion of these protein variants to the truncated RNA polymerase (Figure 5A,B). This rescue also correlated with the compromised ability of these proteins to function as transcription factors when fused to a DNA binding domain (Kwon et al., 2013).

**Figure 5:**
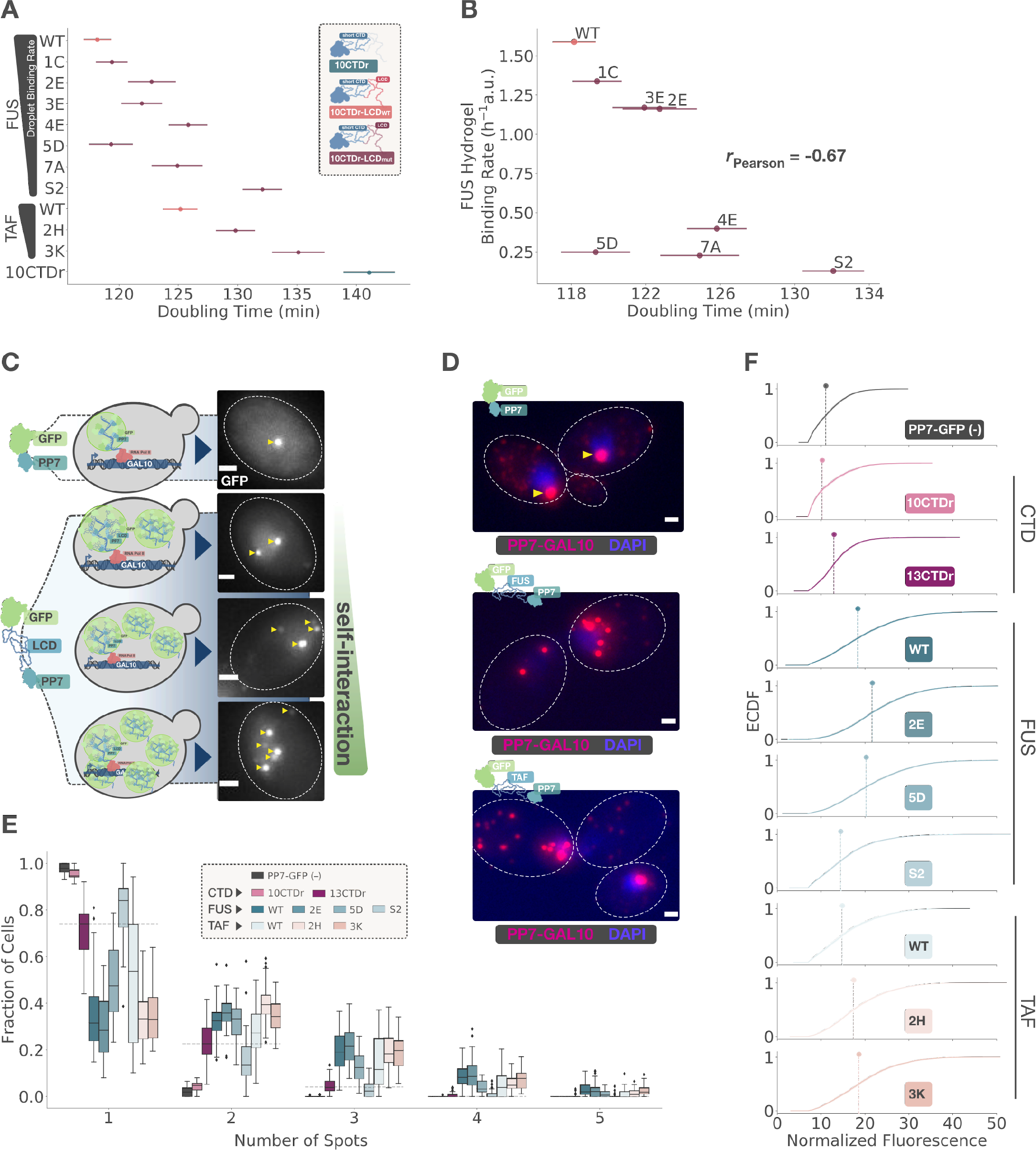
CTD and the low complexity domains (LCD) of FUS and TAF15 can self-interact *in vivo* and this ability is necessary for the function of RNA Pol II. (A) Doubling times (DT) of 10 CTD repeat (CTDr) strains with and without fusion to FUS or TAF15 wild-type and mutated LCDs sorted by droplet binding rate as reported in Kwon et al. (2013) and comparison of FUS mutants with these rates (B). Pearson correlation is indicated. Color indicates whether fused LCD is present, wild-type or mutant. Individual points come from independent lines when available and indicate mean DTs with standard error estimated from three biological replicates per line. (C) Diagram of self-interaction assay. An LCD is fused to GFP and PP7 coat proteins; GFP-PP7 fusion with no LCD is used as control (top). Upon induction of transcription, these fusion proteins bind mRNA scaffolds. If LCDs can self interact, they increase the brightness of spots by recruiting more proteins to the scaffold, and by bringing more than one mRNA together outside of the active transcription site via LCD-LCD interactions (bottom). Representative snapshots of cells with increasing number of GFP fluorescent spots are shown in the right. (D) smFISH after 30 minutes of galactose induction with probes that bind PP7 hairpins in GAL10 mRNA on strains constitutively expressing PP7-GFP with and without LCD fusion as indicated in the upper left corner of each image. Cells without an LCD have a single nuclear RNA complex per cell corresponding to the transcription site (top, yellow arrowheads). Individual mRNA molecules are visible as dimmer spots. Fusion to FUS (middle) or TAF15 (bottom) LCD leads to the formation of multiple RNA complexes, visible as spots brighter than a single mRNA, in individual cells as a result of intermolecular LCD-LCD interactions. White dotted contours mark cell outlines; scale bar is 1*μ*m. (E) Fraction of cells per field o f v iew t hat c ontain e ach n umber o f G FP s pots b y s train f rom t hree b iological r eplicates. The horizontal blue dotted lines indicate the number of spots in the 13CTDr GFP-PP7 fusion. Protein fused to PP7-GFP is indicated by color. (F) Corresponding empirical cumulative distribution functions (ECDF) of GFP fluorescence intensities from the brightest spot in each cell, typically corresponding to the transcription site. Protein fused to PP7-GFP is indicated by color and in the lower right of each plot.

We investigated these mutants more closely by using an assay designed to measure self-interactions *in vivo*, which we define as the ability of a protein to interact with and recruit others of its k ind. We speculated that fusing a self-interacting protein to PP7-GFP would lead to brighter spots in our live transcription assay (Figure 5C, left). Heterologously expressed FUS and TAF15 can form punctate structures in yeast (Couthouis et al., 2011; Ju et al., 2011). However, in this assay wild-type FUS and TAF15 LCD fusions distributed uniformly across the cell nucleus, presumably due to lower protein concentrations. On the other hand, the spots that formed after galactose induction became brighter with either LCD compared to PP7-GFP alone. Moreover, more than a single spot became visible soon after transcription activation (Figure 5C, right).

We characterized this phenomenon via smFISH of galactose induced strains carrying PP7-GFP with and without FUS and TAF15 LCD fusion (Figure 5D). This experiment suggested that the increase in the number of spots and their brightness could be a result of 1) indirect recruitment of PP7-GFP-LCD to each mRNA scaffold but mostly 2) LCD-mediated physical interactions between mRNA molecules outside of the TS. We proceeded to use this assay to determine whether a protein can self-interact at physiological concentrations *in vivo*.

We counted the number of bright spots per cell arising 30 minutes after galactose induction in snapshots taken during a 20 minute window with a maximum intensity laser. This number is almost always 1 for PP7-GFP alone, corresponding to the TS, except for when a cell had just duplicated the GAL10 locus during cell division. A 10CTDr-PP7-GFP protein fusion mostly produced a single spot, similar to the non-self-interacting control. On the other hand, a 13CTDr and all of the FUS and TAF15 variants resulted in a higher fraction of cells with more than a single bright spot (Figure 5E).

We also compared the fluorescence intensity of the brightest spot per cell, presumably the TS, to the control expressing PP7-GFP only. Proteins that formed multiple bright spots also increased the intensity of the brightest spot (Figure 5F). The 10CTDr construct resembled the negative control more than 13 CTDr. Generally, these measurements support that 13CTDr, FUS, and TAF LCDs can self-interact, and the extent of self-interaction qualitatively recapitulates the transitions observed in our growth and transcription assays.

These results suggest that self-interactions are necessary for the transcriptional rescue of CTD truncation, and support the idea that this ability is a key attribute that the CTD’s long disorder contributes to transcription.

### 2.5. An integrative transcription model explains the influence of CTD length

In light of our evidence, we sought to devise a quantitative model for transcription that captured the effects of perturbing CTD length and illuminated the role of disorder-mediated self-interactions. We built upon a model that includes an active and an inactive state, which enables it to produce transcriptional bursting, and specifies that an RNA polymerase molecule can only be recruited during the active state, in agreement with experimental data (Bartman et al., 2019).

The CTD is known to physically interact with Mediator (Thompson et al., 1993; Kim et al., 1994), a prevalent component of the preinitiation complex (PIC) (Allen and Taatjes, 2015). Given the repetitive nature of the CTD, we postulated that the number of repeats and hence CTD length could modulate the affinity with which the polymerase binds a Mediator-bearing PIC. Following this logic, we chose to explicitly refer to the active state as the assembled PIC, primed for RNA Pol II binding.

Polymerase release from the PIC is preceded by CTD phosphorylation (Payne et al., 1989; Svejstrup et al., 1997), which disrupts their physical interaction (Jeronimo and Robert, 2014; Wong et al., 2014). Assuming each of the CTD repeats contributes to this interaction, we reasoned that their number should correlate with the rate of polymerase release. Our rationale is that given the ratio of unphosphorylated to phosphorylated repeats determines the physicochemical state of the CTD and its interaction with Mediator (Robinson et al., 2016), the number of phosphorylation sites, or CTD length, should be proportional to the time it takes for CTD kinases to reach this threshold.

To recapitulate, we postulated that CTD length influences transcription by enhancing polymerase recruitment rate to the PIC (*β*) and by decreasing polymerase release rate (*ϕ*) from the PIC (Figure 6A, blue).

**Figure 6:**
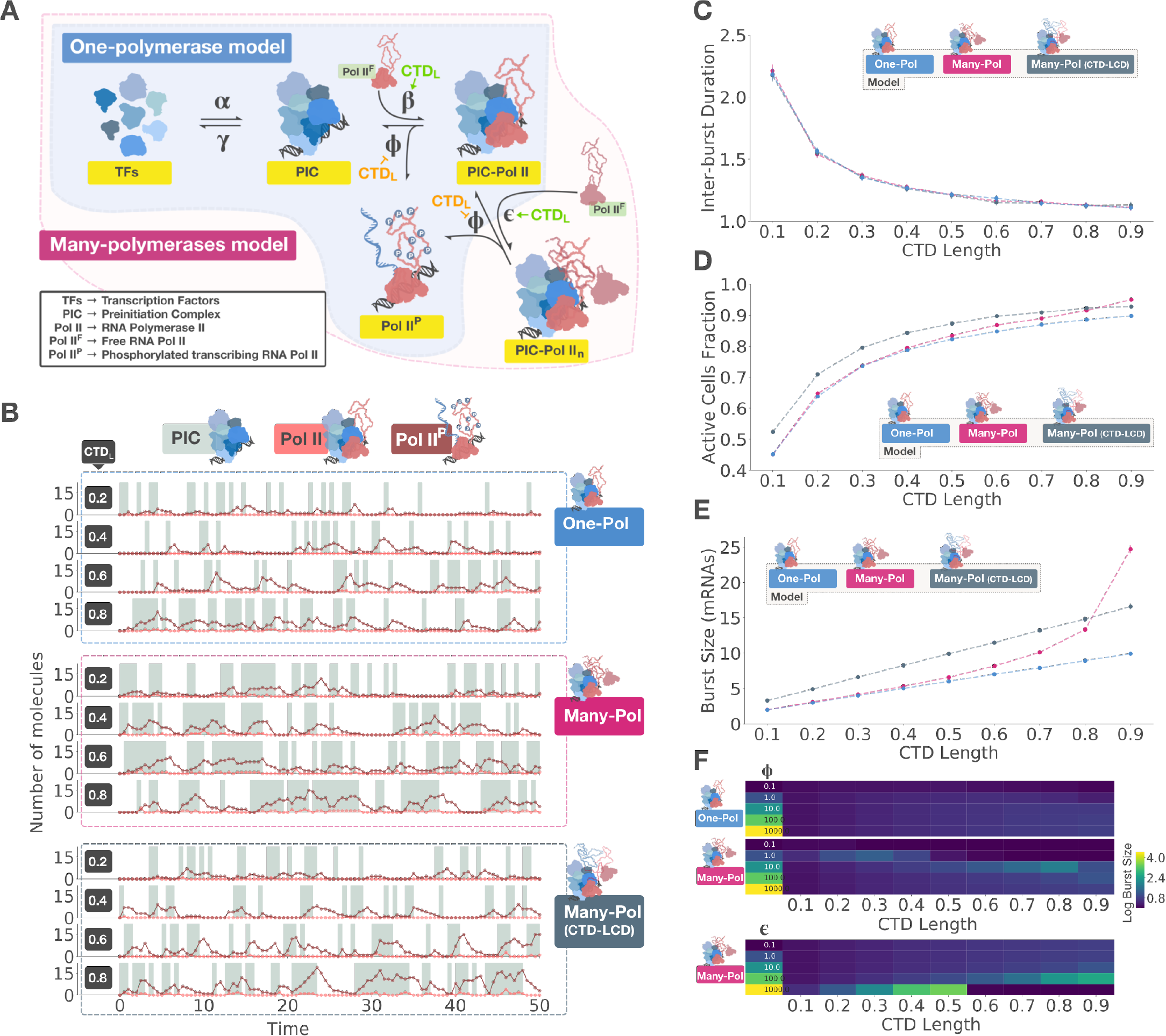
An integrative model for transcription activation explains the role of CTD length. (A) CTD-centric model for transcription activation. Transcription factors (TFs) assemble to form a preinitiation complex (PIC) to which an RNA polymerase binds. In the one-polymerase model (blue), only a single polymerase is allowed to bind the complex at a time. In the many-polymerases model (pink), more than one polymerase can bind the complex via CTD-CTD interactions. Fusion of a truncated CTD to a self-interacting protein is simulated using the many-polymerases model with fixed *ϵ* and *ϕ*. The positive or negative influence of CTD length (*CT D*_*L*_) on each rate is indicated in green and orange, respectively. (B) Representative traces from stochastic simulations by model, indicated to the right, as a function of *CT D*_*L*_, indicated in the left side of each plot. Background shows PIC assembly state; number of PIC bound and phosphorylated (transcribing) polymerases are shown in color as indicated in the legend on top. Corresponding mean inter-burst durations (C), mean fractions of active cells (D) and mean burst sizes (E) with 99% bootstrapped confidence interval. (F) Effect of varying *ϕ* (top) and *ϵ* (bottom) on the log burst size.

An important feature of the current model for transcription is that only a single polymerase molecule can bind the PIC at a given time (Bartman et al. (2019); Figure 6A, blue). To incorporate the ability of the CTD to self-interact, we postulated an additional molecular state that allows for the recruitment of more than single polymerase molecule to the extant PIC-Pol II complex via CTD-CTD self-interactions (Figure 6A, pink).

Finally, we used this model to assess the transcriptional rescue observed upon fusion of FUS or TAF15 LCDs to a CTD-truncated RNA polymerase. We reasoned that these LCDs would contribute the ability to self-interact, but make it more difficult for the polymerase to be released into the gene because their phosphorylation may not be as efficient. Specifically, we postulated that fusing a self-interacting LCD to a truncated CTD would fix self-recruitment rate *ϵ* and release rate *ϕ*, while the rate of initial recruitment *β* would still be determined by CTD length.

We interrogated the consistency of our models with experimental data using stochastic simulations. For simplicity, we set CTD length (*CT D*_*L*_) to be a number in the range (0-1), directly proportional to both polymerase recruitment rates *β* and *ϵ*, whose complement (1 − *CTD*_*L*_) is proportional to polymerase release rate *ϕ*. We visualized these simulations as transcriptional traces for each model, including states of PIC assembly, numbers of PIC-bound and phosphorylated polymerases (Figure 6B and Figure S6), akin to our live transcription imaging data (Figure 3B). From these simulations we computed the distributions of burst sizes, inter-burst times and fraction of active cells, and asked how CTD length would affect these parameters. Importantly, to be consistent with prior literature, we define the start and end of a burst to be concomitant with PIC assembly and disassembly, respectively, and only consider those that yield at least one mRNA molecule.

We compared the outcomes of the one-polymerase, the many-polymerases and the rescue models. The identity of the distributions of burst sizes and inter-burst times resembled geometric and exponential, respectively, for all the models (Figure S7). In addition, because the difference between models lies in the states after the start of a burst and hence does not influence burst frequency, the model choice did not affect the resulting distributions of inter-burst times (Figure 6C, S7C).

The three models predicted that shortening the CTD would progressively increase the average inter-burst time (Figure 6C). This in turn translated into a progressive decrease in the fraction of active cells (Figure 6D). Importantly, the magnitude of change in these numbers increased with decreasing CTD length. This effect qualitatively reproduced the observed transitions in growth and transcription phenotypes resulting from CTD truncation (Figure 2A, 3E).

Although dominated by burst frequency in this regime, the fraction of active cells was also influenced by burst size (Figure S8A). Our simulations predicted that such influence would lead to a modest but consistent increase in the fraction of active cells across a range of CTD lengths when comparing rescued with truncated CTD lengths (Figure 6D). In contrast to frequency, burst size was significantly influenced by model choice. The rescue model predicted a consistently higher mean burst size at almost every CTD length, except at the extreme of a long CTD (Figure 6E). These results are strikingly consistent with our experiments, in that rescued strains show a moderate increase in the fraction of active cells compared to truncated CTDs (Figure 4K,L), and TS intensity increased upon LCD fusion (Figure 4M,N). These comparisons of simulations with experiments support the existence of a state with more than one polymerase and that LCD fusion specifically rescues burst size via self-interactions.

The one-polymerase and the many-polymerases models produced very similar mean burst sizes with short CTDs. However, burst sizes resulting from each model deviated significantly with longer CTDs (Figure 6E). In particular, they increased exponentially when many polymerases were allowed to bind the PIC, but only linearly with a single polymerase. These predictions suggest the impact of binding more than one polymerase may become increasingly relevant as the CTD grows longer.

We also sought to elucidate the relationship between burst size, duration, and TS intensity in our experiments by comparing these three values in our simulations. Because CTD length inhibits release rate in the one-polymerase and the many-polymerases models, burst duration increased faster than burst size (Figure S8B); these parameters were linearly proportional only when release rate was fixed in the case of the rescue model. These results potentially explain why burst duration decays more subtly (Figure S3B-D) than TS intensity (Figure 3D,F). A potential source of uncertainty in measuring burst size is that the decay in TS intensity could be a result of decreased burst frequency, given that GAL10 is transcribed in highly frequent bursts that could overlap in time and inflate the measured intensity of a burst. We simulated how this intensity signal would vary across CTD lengths and under different elongation rates *δ*, which determines how long a given mRNA stays and contributes to the TS signal. We compared the fraction of active cells with the difference between this observed TS intensity, influenced by *δ* and measured as the distributions of peak intensities in the final simulated traces, and the true burst size, which we kept track of as we generated the traces and is independent of *δ* (Figure S8C). The trend in this analysis is that if *δ* is low, the fraction of active cells would remain high across CTD lengths and TS intensity would overestimate the true burst size because sequential bursts would overlap; if *δ* is large, the fraction of active cells would remain low, and TS intensity would underestimate the true burst size because intensity would decrease before the end of a burst. Both of these effects were enhanced with longer CTDs. The experimental range of active cells fractions (Figure 3E) suggests a scenario where the estimate from TS intensity lies between a slight overestimation to an underestimation of the true burst size; in the latter situation, inferring burst size from these data would be a conservative estimate. In the context of the canonical transcription model (Figure 6A), the polymerase binding rate *β* intrinsically links burst size and frequency. We thus conclude that the simplest explanation consistent with simulations, experiments and previous literature is that burst size and frequency both decrease with CTD truncation.

Varying each of the model parameters individually (Figure S8E,F) offered an additional insight. By modulating the rate of polymerase phosphorylation *ϕ*–conceivably in local nuclear environments that limit CTD kinase activity– or by increasing the rate of self-interaction *ϵ*, it is possible to dramatically increase burst size under the many-polymerases model (Figure 6F). This effect would be a direct consequence of increased concentrations of unphosphorylated Pol II, akin to recently reported droplets in cells (Chong et al., 2018; Boehning et al., 2018; Cho et al., 2018; Nair et al., 2019). Although not specified in our model, liquid-liquid phase separation may thus naturally emerge from this transcription logic.

Our CTD-centric models helped us understand our experimental observations and integrate them with prior knowledge. By simulating them, we were able to capture the behavior of transcription upon CTD truncation and subsequent fusion to LCDs, illuminating the role of CTD length and providing support for a novel molecular state where more than a single polymerase can bind the PIC.

## 3. Discussion

### 3.1. A CTD-centric model offers mechanistic insights into transcriptional bursting

RNA Polymerase II is essential and extremely conserved across eukaryotes, yet the amino acid length of its catalytic subunit varies dramatically (Figure 1A,B). As discussed above, the number of disordered amino acids in the CTD closely follows this length variation, increasing with genome size as coding sequences become more scattered (Figure 1C,D).

The influence of CTD length on transcription is consistent with a simple quantitative model based on known protein-protein interactions with Mediator, CTD phosphorylation and disorder-mediated self-interactions (Figure 6A). We explicitly assume that at this level of abstraction, the major, Poissonian source of stochasticity in transcription comes from the multimolecular assembly of the preinitiation complex (PIC) at the enhancer and the promoter, ocurring at the start of each burst and preceding Pol II recruitment. This assumption is motivated by the CTD’s influence on both burst size and frequency (Figure 3C-F) and by extensive previous experimental evidence. PIC formation is a rate-limiting step in transcription (Kuras and Struhl, 1999; Li et al., 1999) and transcription bursts are concomitant with enhancer-promoter interactions (Bartman et al., 2016; Chen et al., 2018). Roughly, the likely order of protein recruitment events upon activation is enhancer specific transcription factors, a single Mediator complex (Petrenko et al., 2016) and general transcription factors, and finally RNA polymerase followed by CTD kinases (Bryant and Ptashne, 2003; Krishnamurthy and Hampsey, 2009). In specifying our model, we put forward a view in which bursts are a result of recurrent Pol II binding to the assembled PIC, and inactivity periods a consequence of PIC disassembly (Figure 6A). This reductionist framework thus offers an intelligible perspective of the mechanism of eukaryotic transcriptional bursting.

### 3.2. CTD length cooperatively scales transcription

The probability of interaction between two genomic loci is highly dependent on the physical distance between them (Lieberman-Aiden et al., 2009). As a result, order-of-magnitude variation in genome sizes and in the physical spacing between enhancers and promoters represents a challenge: how does the transcription machinery overcome an increasingly infrequent event? Our simulations suggest increasing CTD length can reduce the number of times an assembled PIC fails to recruit RNA Pol II and produce mRNA before disassembly (Figure S8E), and that self-interactions can considerably increase burst size (Figure 6E,F). By exploiting each rare assembly event, CTD length-enhanced recruitment and self-interactions could contribute to resolve the transcription scaling paradox. This compensatory mechanism would complement changes in genome organization (Szabo et al., 2019), without which enhancer-promoter interactions may never occur in the first place.

In this context, CTD length bears an important distinction with the strength of self-interaction and polymerase recruitment rate, determined by interactions with the PIC. While the naive expectation is that these parameters should be correlated, devations from the consensus repeat YSPTSPS may provide a way to modulate them independently. This hypothesis could explain why fruit flies with a CTD o f wild-type length that is made up entirely of consensus repeats do not survive, but animals with a yeast CTD remain viable (Lu et al., 2019). Based on this rationale, we speculate CTD length and sequence in eukaryotes coevolves with the physical spacing between enhancers and promoters, primarily determined by genome organization (Lieberman-Aiden et al., 2009; Szabo et al., 2019) and genome size.

### 3.3. CTD-CTD self-interactions link transcription activation to phase-separation

It is possible that FUS and TAF15 LCDs rescue CTD truncation through an alternative recruitment mechanism that does not involve self-interactions, given they can also function as transcription factors when fused to a DNA binding domain (Kwon et al., 2013). This hypothesis would nonetheless be consistent with our inference that CTD length modulates polymerase binding rate to the PIC, but it would not support a many-polymerases state. On the other hand, our data do not suggest rescue is driven by enhanced direct recruitment to the PIC, given the increase in fraction of active cells seems to be predominantly driven by burst size and not frequency (Figure 4K-N), while direct recruitment would enhance both parameters. Additionally, we find a correlation between LCD ability to bind liquid droplets (Figure S4D) and self-interact with the extent of phenotypic rescue upon fusing them to a truncated RNA Pol II (Figure 5A,C,D).

Self-interactions additionally offer a logical connection between the mechanism of transcription activation and liquid-liquid phase separation (LLPS). Our model predicts that when self-interaction strength is large or the rate of polymerase release is small, large transcription bursts could emerge (Figure 6F), implying a high local concentration of unphosphorylated polymerases. This environment has been observed in LLPS droplets at super-enhancers of live cells (Chong et al., 2018; Boehning et al., 2018; Cho et al., 2018; Nair et al., 2019). A corollary of this idea is that the average gene and the super-enhancer gene can both be transcribed using the same mechanisms, but only the latter would manifest LLPS droplets as an epiphenomenon of enhanced polymerase recruitment or kinase exclusion. Super-enhancers would then be at the extreme of the distribution of burst sizes, which is consistent with the observation of only a few droplets per cell whose number does not nearly match the total number of transcribed genes. In this scenario, LLPS could result in emergent behaviors whose understanding would require a different quantitave framework; our model may not apply to these CTD-lengths but could provide a useful expectation to compare them with. In other words, CTD length variation may result in regimes of transcription activation governed by different dynamics.

### 3.4. Self-interactions support a multi-polymerase complex

The key proposition of our model that allows the incorporation of self-interactions is the existence of a molecular complex that can bind more than one RNA Polymerase molecule (Figure 6A, pink). Short-lived Pol II clusters observed in mammalian cells that overlap with active transcription sites and whose duration correlates with mRNA output (Cisse et al., 2013; Cho et al., 2016) could be a direct observation of this event. On the other hand, Pol II pausing appears to negatively correlate with transcription initiation (Shao and Zeitlinger, 2017; Gressel et al., 2017), which could suggest that new polymerases may not be able to bind an occupied promoter. Distinguishing the perhaps differential ability of PIC-bound and paused Pol II’s CTD to self-interact would be helpful to understand the relationship of this observation with a many-polymerases state.

Pol II is released from the promoter upon CTD phosphorylation (Jeronimo and Robert, 2014; Wong et al., 2014), based on which we argue that CTD length influences release rate. Along this line, depletion of yeast CTD-kinase Kin28 causes an upstream shift in Pol II occupancy along genes (Wong et al., 2014), with a pattern that resembles proximal-promoter accumulation in metazoans (Adelman and Lis, 2012) and is consistent with a defective promoter escape. A conspicuously similar shift was observed upon mutating CTD’s serine 5 (Collin et al., 2019), the specific CTD residue phosphorylated for transcription initiation (Eick and Geyer, 2013; Harlen and Churchman, 2017). We find self-interactions correlate with the efficiency of transcription (Figure 5). A sensible interpretation of these experiments is that decreasing CTD-kinases or their activity on the CTD lead to increased RNA Pol II at the promoter by extending the time window for self-interaction mediated recruitment. These observations raise the hypothesis that promoter accumulation of Pol II in metazoans (Adelman and Lis, 2012), congruently not observed in yeast (Steinmetz et al., 2006), could be contingent on a higher phosphorylation release-threshold linked to a long CTD and a multi-polymerase complex. Experiments that directly measure the number of polymerases that can bind the PIC, and how CTD length influences RNA Pol II occupancy profile would be highly informative in this regard.

In summary, our study integrates experimental results and simulations to explain how CTD length influences transcription activation. We revise the current model of transcription by providing evidence that self-interactions are a key feature in this process, intrinsically linked to a state in which multiple polymerases can bind the PIC. This line of reasoning offers a sound connection between a reductionist, concrete transcriptional logic and the emerging perspective of phase-separation, generating testable hypotheses that will further clarify the functional and evolutionary relevance of CTD length variation.

## 4. Acknowledgements

We thank Mitchell Guttman, Matt Thomson and members of the Sternberg laboratory for helpful discussions, Heun J. Lee, Andres Collazo and Giada Spigolon for imaging assistance, Igor Antoshechkin and Vijaya Kumar for RNA-seq experiments, and Steven Mcknight and Masato Kato for reagents. This work was supported by the Howard Hughes Medical Institute with which PWS was an investigator, by the Gordon Ross Medical Foundation and the Benjamin M. Rosen Graduate Fellowships, by the Biological Imaging Center at the Caltech Beckman Institute, and by the Millard and Muriel Jacobs Genetics and Genomics Laboratory.

## 5. Author Contributions

Conceptualization, P.Q.C., P.W.S; Methodology, P.Q.C, P.W.S., T.L.L.; Software, P.Q.C.; Formal analysis, P.Q.C., T.L.L.; Investigation, P.Q.C.; Data Curation, P.Q.C.; Writing – Original Draft, P.Q.C., P.W.S.; Writing – Review & Editing, P.Q.C., P.W.S., T.L.L.; Visualization, P.Q.C.; Supervision, P.W.S.; Funding Acquisition, P.W.S.

## 7. Methods

### 7.1. Data analysis

Except when indicated, all programming, data extraction, wrangling, calculations and plotting were done using Python 3.7 with standard scientific libraries (Oliphant, 2007; Jones et al.; Millman and Aivazis, 2011). All scripts used in this paper are available in the following github repository: https://github.com/WormLabCaltech

### 7.2. Image analysis

Maximum-intensity projections were used for all z-stack images, sometimes generated and often visualized using Fiji (Schindelin et al., 2012).

Cells were segmented using local thresholds and the Watershed algorithm. Candidate 2D fluorescent peaks were detected and tracked using Trackpy (Allan et al., 2018) with minor adaptations.

For PP7 transcription dynamics imaged with low laser intensity, only the brightest peak per cell per frame was kept. A Gaussian-Process Classifier (GPC) trained with a set of manually classified images was then used to distinguish transcription sites from spurious peaks, only keeping those with a GPC probability of at least 0.5 (Figure S2A-C). Transcription intensity was expressed as the fold-change of peak over mean nuclear fluorescence. This metric yielded overlapping intensity distributions of the same strain imaged with different settings (Figure S2D-H). Autocorrelation analysis was carried out as previously described (Lenstra and Larson, 2016). Missing timepoints where no peak was detected were imputed using the intensity at the position of the previous spot. For snapshots and smFISH images taken with maximum laser intensity, in which signal-to-noise ratio was greater, manually determined intensity thresholds were used.

### 7.3. RPB1 bioinformatic analysis

RPB1 homologs were retrieved by searching Ensembl database (Zerbino et al., 2018) using the HMMER online tool (Finn et al., 2011) with default settings, starting with the yeast RPB1 protein sequence. Amino acid sequences were analyzed for disorder locally using MobiDB-lite (Necci et al., 2017), which provides a consensus score derived from eight disorder predictors, in a machine running Unix Debian 4.9 and Python 2.7. Genome sizes and gene numbers were scraped from Ensembl websites using a custom script.

### 7.4. Genetic constructs

All constructs used in this paper were built using PCR amplification, Gibson (Gibson et al., 2009) or golden gate (Engler et al., 2008) assembly methods and verified by Sanger sequencing. Plasmids are listed in Table S1.

Wild-type LCDs of FUS (residues 1-214) and TAF15 (residues 1-208) were as previously defined (Kwon et al., 2013) and obtained by PCR amplification from human cell line 293T cDNA. Plasmids with coding sequences of previously reported FUS and TAF15 LCD tyrosine-to-serine mutants (Kwon et al., 2013) were a gift from Steven Mcknight.

The DNA sequence for CTD truncation repair templates was redesigned to facilitate PCR amplification, and together with yeast codon-optimized mScarlet coding sequence, synthesized as an Integrated DNA Technologies (IDT) gBlock and cloned into their respective vectors. sgRNAs were purchased as individual oligos, hybridized and cloned into pWS082 using golden gate assembly.

### 7.5. Strain Engineering

All transformations were carried out using the LiAc/SS Carrier DNA/PEG method (Daniel Gietz and Woods, 2002). Strains are listed in Table S2.

Strain YTL047A (Donovan et al., 2019) was generated by transforming diploid *S. cerevisiae* BY4743 with a PCR product containing the PP7 loop cassette and a loxP-kanMX-loxP marker, which was subsequently removed with Cre recombinase. A single allele of GAL10 was tagged. All strains used in this study are derivatives of YTL047A.

RPB1 modifications were engineered in both alleles using CRISPR-Cas9 with gRNAs of improved stability (Ryan et al., 2014), antibiotic-mediated selection of cells proficient in gap repair (Horwitz et al., 2015), and plamids from the Yeast MoClo Toolkit (Lee et al., 2015) modified by Tom Ellis’s lab. sgRNA sequences are listed in Table S3.

CTD truncations, mScarlet and LCD RPB1 strains were generated by transforming YTL047A with 100 ng of BsmBI linearized and gel-purified Cas9-kanR plasmid (pWS173), 200 ng of each EcoRV linearized sgRNA vector (pWS082 derivatives), and 2-5 ug linearized repair template, selected for with G418. Correctly modified strains were identified using PCR of zymolyase digested colony scrapes followed by Sanger sequencing.

Strains for live transcription imaging were generated by integrating a single copy of GFP-Envy (Slubowski et al., 2015) fused to PP7 coat protein under an rpl15A promoter and a functional ura3 gene into the ura3Δ0 locus by transforming PacI linearized pTL174 and selected for with plates lacking uracil. Strains for self-recruitment assays (Figure 5) were constructed in the same way, except integrating PP7-LCD-GFP fusions. smFISH of self-recruitment assays was done on YTL047A transformed with plasmids pTL092, pQC075 or pQC076 (Figure 5D).

### 7.6. Cell growth measurements

Optical density (OD) was measured at an absorbance wavelength of 600 nm for 16 to 24 hours every 15 minutes using 1:100 dilutions of overnight cultures in 150 uL of YPD in a Falcon flat-bottom 96-Well Clear Assay Plate with lid on a Biotek Cytation 3 microplate reader with 1000 rpm shaking at 30C.

Doubling times were estimated non-parametrically from time derivatives of OD measurements with Gaussian processes using previously described software (Swain et al., 2016).

### 7.7. Live fluorescence microscopy

All microscopy experiments were done using early to mid-log cultures (typically 5e6 to 1e7 cells/mL) growing at 30C with 250 RPM shaking.

mScarlet-RPB1 strains were imaged on 2% agarose pads on coverslips at room temperature immediatly after spinning down at 3600 RCF cultures growing in Synthetic Complete (SC) 2% Glucose media on a Zeiss Imager Z2 microscope with an Axiocam 506 Mono camera, 63x oil objective, 150ms exposure time and 25% laser intensity.

Live transcription and self-interaction imaging was done on concanavalin-A-coated MatTek dishes at 30C as previously described (Lenstra and Larson, 2016) using an Leica DMI 6000 wide-field fluorescence microscope with an Andor Zyla 5.5 or a Hamamatsu Flash 4.0 v3 camera with a 100x oil objective. Cells were induced by adding galactose dissolved in 2 mL SC to 1 mL SC 2% raffinose for a final 3 mL SC 2% galactose and imaged immediately every 20 sec for around 1 hour (live transcription) or after 30 min for 20 min (self-interaction). Live movies were taken with 150 ms exposure, 9 z-stacks every 0.5 *μ*m, and minimal laser intensity to avoid photo-toxicity. Snapshots were imaged once per field-of-view with maximum laser power, 150 ms exposure, and 9-15 manually set z-stacks every 0.5 *μ*m.

### 7.8. RNA-seq

RNA was extracted from mid-log cultures growing in SC 2% raffinose after 2h of 2% galactose or blank induction using Zymo Quick-RNA Fungal/Bacterial Microprep Kit (Catalog # R2010) lysed in an MP Biomedicals FastPrep-24 machine.

RNA integrity was assessed using RNA 6000 Pico Kit for Bioanalyzer (Agilent Technologies #5067-1513) and mRNA was isolated using NEBNext Poly(A) mRNA Magnetic Isolation Module (NEB #E7490). RNA-seq libraries were constructed using NEBNext Ultra II RNA Library Prep Kit for Illumina (NEB #E7770) following manufacturer’s instructions. Briefly, mRNA isolated f rom 1 *μ*g of total RNA was fragmented to the average size of 200 nt by incubating at 94C for 15 min in first s trand b uffer, c DNA w as synthesized using random primers and ProtoScript II Reverse Transcriptase followed by second strand synthesis using NEB Second Strand Synthesis Enzyme Mix. Resulting DNA fragments were end-repaired, dA tailed and ligated to NEBNext hairpin adaptors (NEB #E7335). After ligation, adaptors were converted to the ‘Y’ shape by treating with USER enzyme and DNA fragments were size selected using Agencourt AMPure XP beads (Beckman Coulter #A63880) to generate fragment sizes between 250 and 350 bp. Adaptor-ligated DNA was PCR amplified followed by AMPure XP bead clean u p. Libraries were quantified with Qubit dsDNA HS Kit (ThermoFisher Scientific #Q32854) and the size distribution was confirmed with High Sensitivity DNA Kit for Bioanalyzer (Agilent Technologies #5067-4626). Libraries were sequenced on Illumina HiSeq2500 in single read mode with the read length of 50 nt following manufacturer’s instructions. Base calls were performed with RTA 1.13.48.0 followed by conversion to FASTQ with bcl2fastq 1.8.4.

RNA-seq quantification was performed using Kallisto (Bray et al., 2016) with 200 bootstraps in sigle-end mode with average length of 300 bp and standard deviation of 20 bp. Differential expression analysis was done with Sleuth (Pimentel et al., 2017) using the linear models described in the results and supplementary sections.

### 7.9. smFISH

smFISH experiments were carried out as described previously (Trcek et al., 2012; Lenstra et al., 2015). TYE665 labeled PP7 probes were purchased from IDT. A set of 48 Quasar 570 labeled probes were designed to target the coding sequence of Gal3 and purchased from Biosearch Technologies. Probe sequences are listed in Table S3.

Mid-log yeast cultures were fixed with paraformaldehyde and permeabilized with lyticase. Hybridization solution with 0.1 uM probes, 10% dextran sulfate, 10% formamide, and 2x Sodium Saline Citrate (SSC) was used to hybridize probes in fixed cells for 4 hours at 37 C. Coverslips were washed twice for 30 min with 10% formamide, 2x SSC at 37C, followed by rinses with 2x SSC, and 1x PBS for 5 minutes. PLL-coated 18 mm diameter #1.5 thickness coverslips were purchased from Neuvitro, mounted on microscope slides using ProLong Gold or Glass Antifade Mountant with DAPI (Life Technologies).

smFISH samples were imaged at room temperature with maximum laser power, 300 ms exposure and 9-15 manually set z-stacks every 0.2 *μ*m on the Leica microscope with 100x objective described above.

### 7.10. Stochastic simulations

Stochastic simulations were performed using software described in Bois and Elowitz (2019) with minor modifications to extract burst start, end and size while generating Gillespie samples. Rates were chosen according to Bartman et al. (2019), with *α* = 1, *γ* = 3, *β* = 30, *ϵ* = 10 and *ϕ* = 100. For trace visualization purposes, a rate of phosphorylated Pol II removal (elongation rate) *δ* = 1 was used.

---

**Figure S1:**
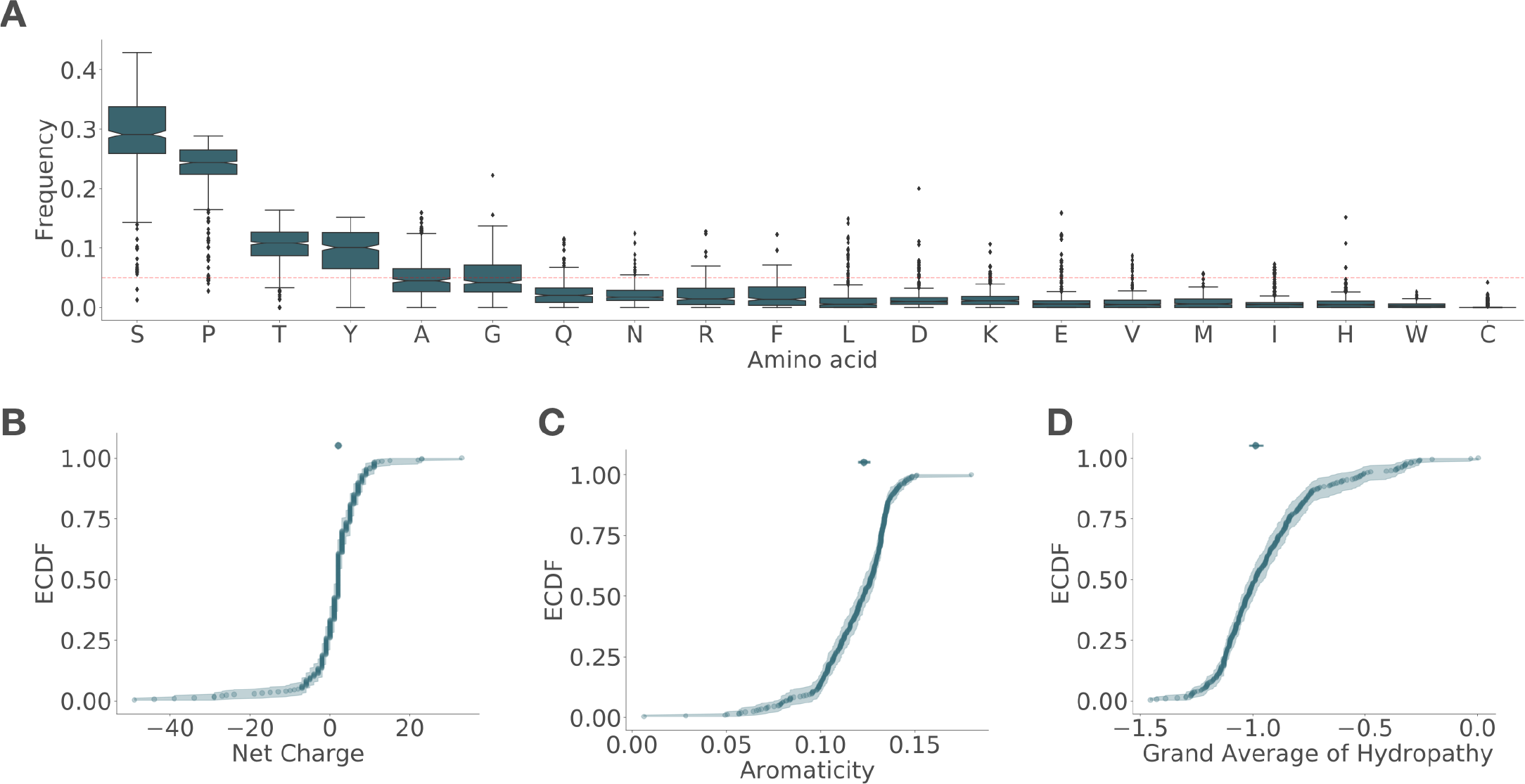
CTDs share amino acid composition. Related to Figure 1. CTDs were identified as the longest contiguous disordered region in RPB1 sequences. Only the longest protein per genus was considered. (A) Amino acid frequency sorted by mean abundance. Red dotted horizontal line indicates a uniform amino acid frequency of 1/20. Empirical cumulative distributions (ECDF) of net charges (B), aromaticity (C) and hydrophobicities (D) based on the grand average of hydropathy score (Kyte and Doolittle, 1982). Shaded area is bootstrapped 99% confidence interval (CI) and top markers show median with 99% CI.

**Figure S2:**
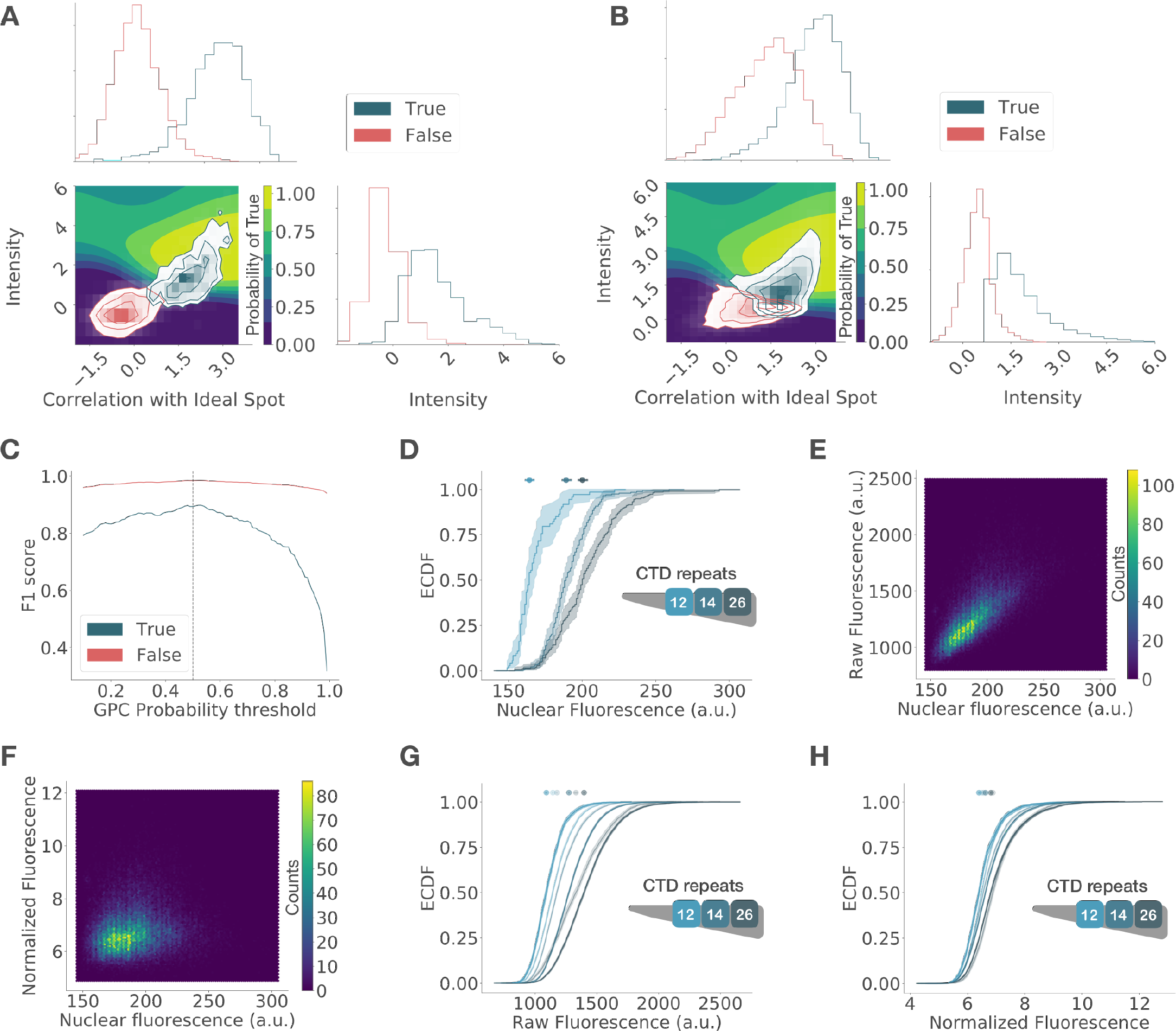
Classification and normalization of PP7-GFP spots enables quantification of transcription dynamics and cross-strain comparisons. Related to figure 3. Candidate spot images were obtained automatically using Trackpy’s (Allan et al., 2018) peak detection algorithm. A sample of this image set was manually classified as True or F alse. For c lassification, spot images were represented using two features: correlation with an ideal spot (a single light point source blurred with a 2D Gaussian function) and intensity. (A) Histograms show the distribution of correlations (top) and intensities (right) of manually labeled spots. Left corner plot shows the joint distributions. This 2D data set was used to train a Gaussian-Process Classifier (GPC), resulting in the decision surface shown underneath, whose color indicates the probability of being a true spot. Candidate spots with a GPC probability above 0.5 were classified as True (B). This threshold was determined based on the change in the accuracy of classification (C), measured using the F1 score on a test set. The vertical dotted line indicates this probability threshold. Mutant strains show different PP7-GFP expression levels, as seen in the empirical cumulative distribution functions (ECDF) of mean nuclear fluorescence by strain (D). These differences result in a correlation observed in the hexagonal bin plot comparing mean nuclear fluorescence with raw spot fluorescence (E), which is removed after normalization (F). Normalized fluorescence is the ratio of spot fluorescence over mean nuclear fluorescence. The efficacy of normalization can also be seen in the ECDFs of raw burst fluorescence by strain imaged with two laser intensities that artificially shift the intensity distributions of the same strains (G), which overlap after normalization (H). Transparency is used to indicate a different laser intensity. Shaded area is bootstrapped 99% confidence interval (CI) and top markers show median with 99% CI.

**Figure S3:**
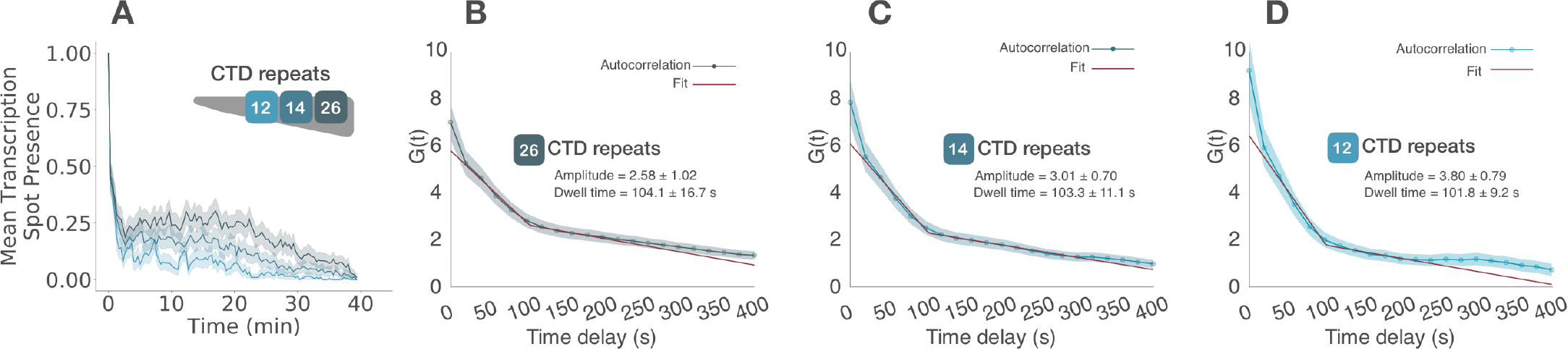
Transcription burst frequency remains constant after activation and decreases with CTD truncation. Related to Figure 3. (A) Mean aligned GAL10-PP7 boolean transcription traces. Boolean traces were obtained by marking with 1 and 0 the presence or absence of a transcription spot (TS), respectively. These traces were aligned and trimmed to begin with the first appearance of a TS and averaged over time, only considering cells that were active during the movie. These traces show the average frequency remains mostly constant over time and decreases with CTD length. Shaded area is bootstrapped 95% mean confidence interval. Frequency decay is also evident from an increase in amplitude, inversely related to frequency, in the autocorrelation of intensity traces corrected for non-steady-state effects in wild-type (B), 14 (C) and 12 (D) CTDr strains. Shaded area indicates standard error of the mean.

**Figure S4:**
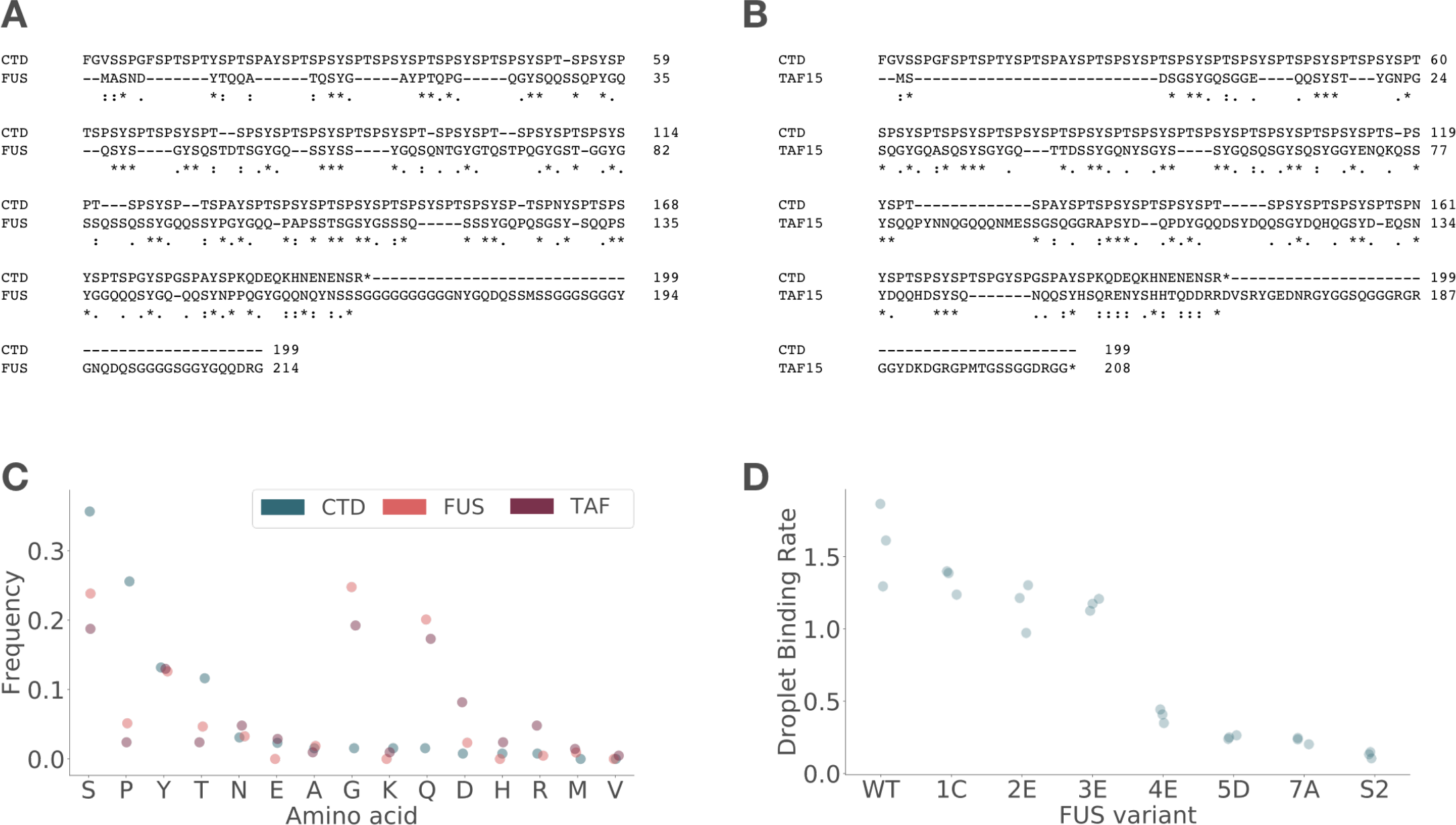
FUS and TAF15 low complexity domains (LCD) are different in sequence but similar in amino acid composition to the CTD. Related to figures 4 and 5. Protein alignments of FUS (A) and TAF15 (B) LCDs with yeast CTD. (C) Amino acid frequency in each of these proteins, sorted by CTD frequency. Only amino acids present in at least one protein are shown. (D) *In vitro* droplet binding rates of FUS variants used in this study. These numbers are the slopes obtained from a linear regression of LCD-GFP binding to wild-type FUS LCD droplets, measured as droplet fluorescence intensity over time. Each point is an experimental replicate; data are from Kwon et al. (2013).

**Figure S5:**
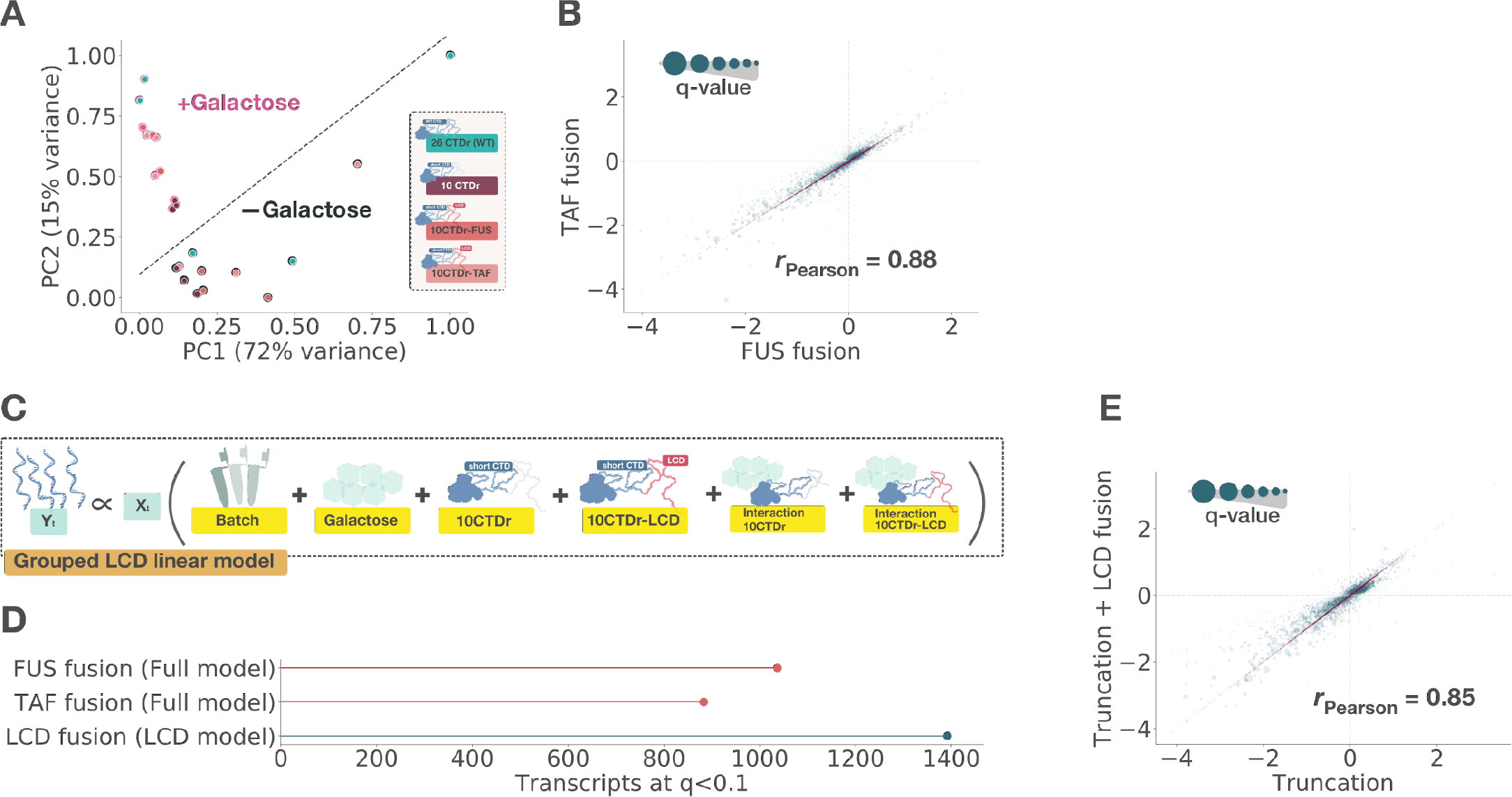
Fusion of a CTD-truncated polymerase to FUS or TAF15 low complexity domains (LCD) results in convergent transcriptomes. Related to figures 2 and 4. (A) Principal compoment analysis (PCA) with the first two PCs scaled to the range [0,1], which together explain 87% of the variance. Each strain has three biological replicates and two conditions. Marker edge color indicates the presence (pink) or absence (black) of galactose in the media; these groups are also divided by the dotted diagonal line. (B) Comparison of the log fold-change of each transcript resulting from FUS and TAF LCD fusion to 10CTDr truncated RNA Pol II under the full linear model shown in Figure 4B. Red points show the positions on the diagonal *x* = *y*. Marker size of each point is inversely proportional to the q-value of the interaction (*ms* = −*log*(*q*_*int*_)); dotted lines reference no change at zero and the Pearson correlation is indicated. (C) Alternative linear model where FUS and TAF rescued strains are grouped together. This grouping results in a higher number of genes identified for LCD fusion under a q-value threshold of 0.1 than for individual coefficients (D). Using this model, (E) comparison of the log fold-change of each transcript resulting from truncation with and without LCD fusion.

**Figure S6:**
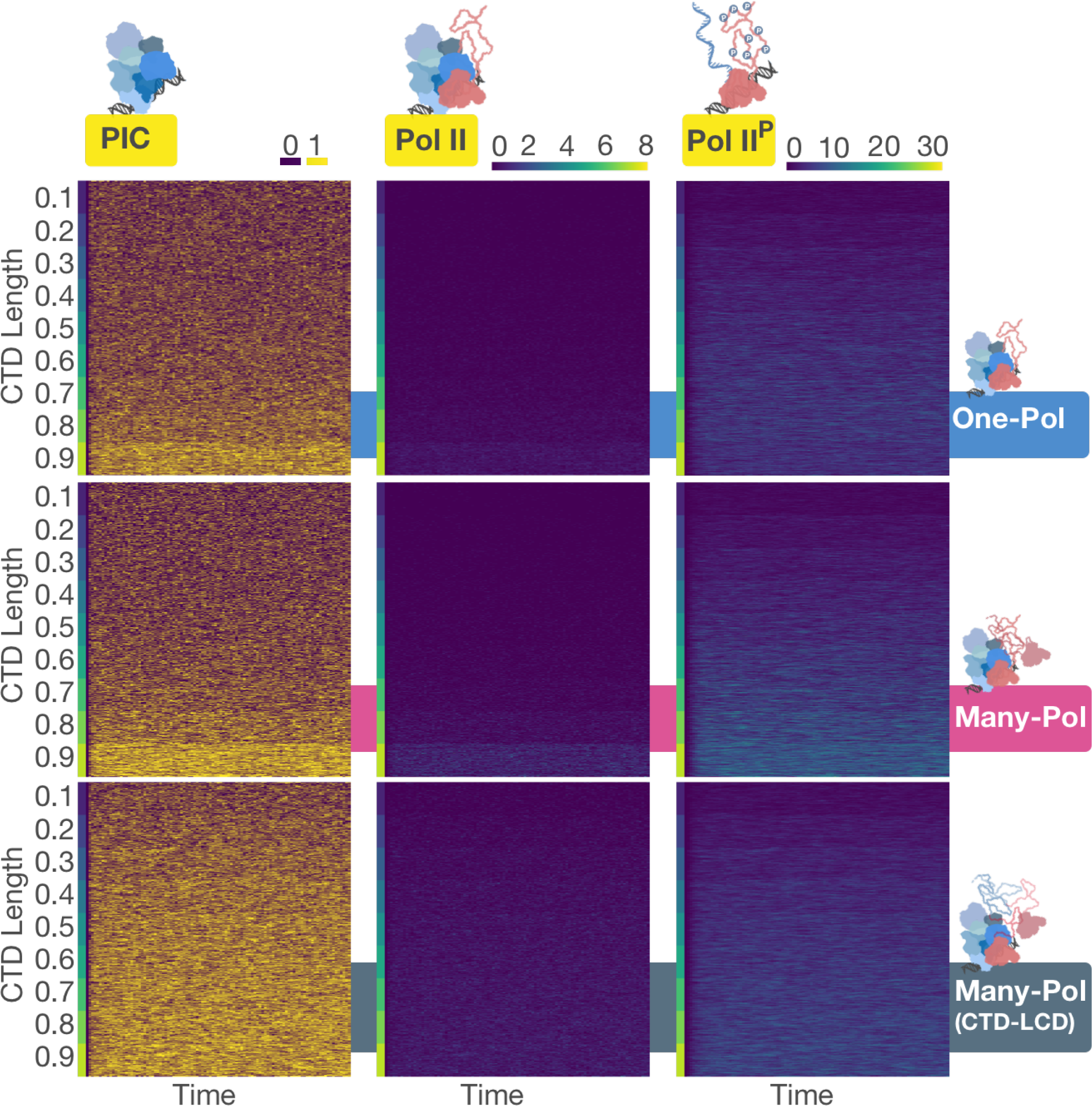
Gillespie simulations yield traces akin to live transcription imaging. Related to figure 6. Traces from stochastic simulations of PIC assembly states, number of PIC bound and phosphorylated (transcribing) polymerases for each model as a function of *CTD*_*L*_, indicated with a colorbar to the left of each panel.

**Figure S7:**
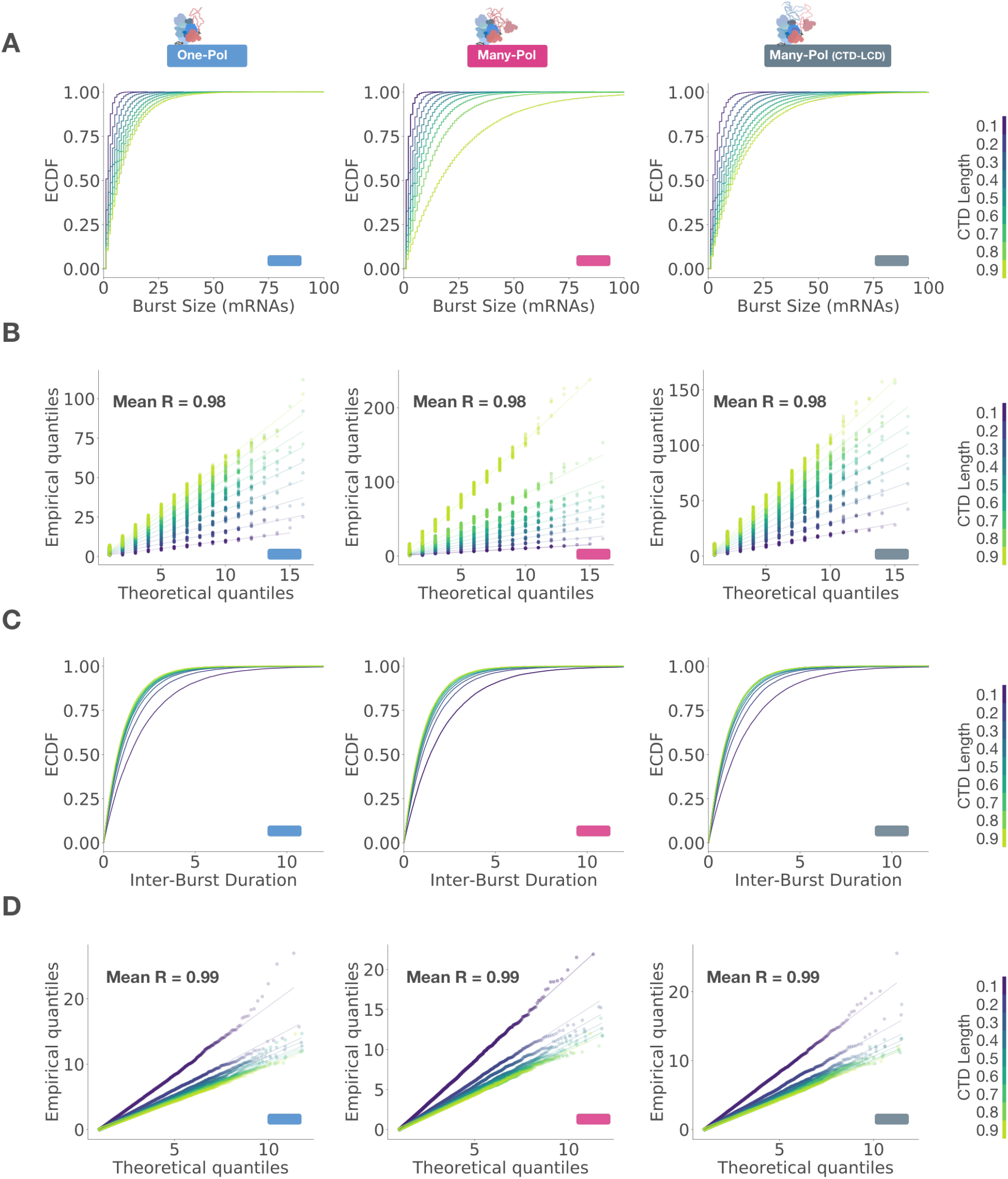
Transcription models produce geometric and exponential distributions of burst sizes and inter-burst durations, respectively. Related to figure 6. (A) Empirical cumulative distribution functions (ECDF) of burst sizes by CTD length for each model. (B) Q-Q plots comparing quantiles from simulated distributions and a geometric distribution. Similarly, (C) ECDFs of inter-burst durations and (D) Q-Q plots comparing their quantiles with an exponential distribution. Each column comes from the model indicated by the color on top and the lower right corner in each plot. The mean of the square root of the coefficient of determination (R) by model is indicated in each quantile comparison. CTD length is indicated by the color shown to the right of each row.

**Figure S8:**
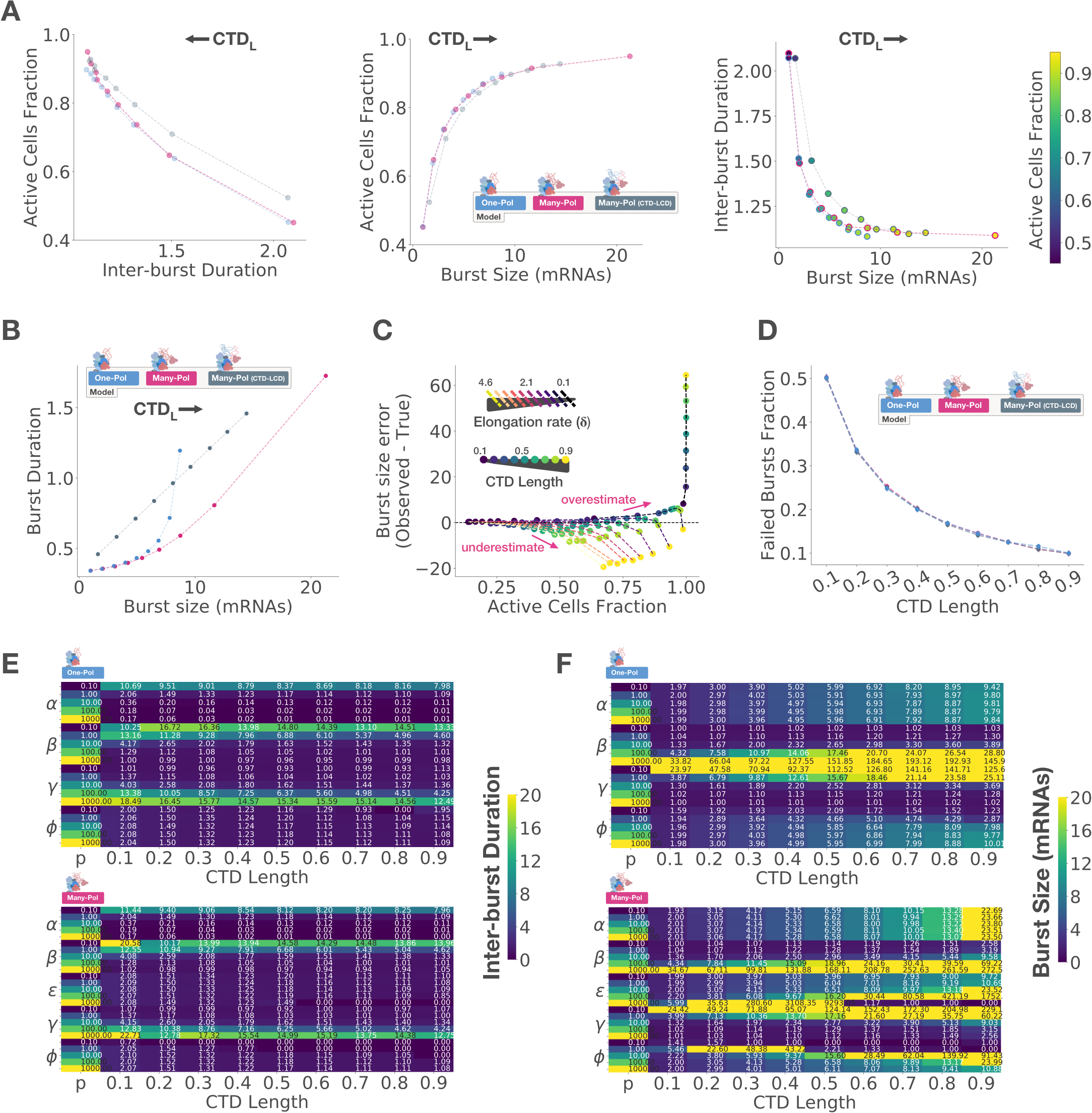
Parameter exploration with stochastic simulations provides insights into experimental observations. Related to figure 6. (A) Comparison of the mean active cells fraction with means of inter-burst duration (left), burst size (middle) and both of these numbers (right) with increasing CTD length (*CTD*_*L*_) by model, indicated with color. Direction of CTD increase is indicated with an arrow on top of each plot. Comparison of mean burst duration with mean burst size with increasing CTD length by model. (C) Comparison of the error in burst size estimate, computed as the difference between the means of the observed transcription site intensity and the true burst size, with the fraction of active cells as a function of *CTD*_*L*_ under the many-polymerases model. The elongation rate (*δ*) determines the time that a given mRNA spends bound to the transcription site and contributes to the observed intensity, thus influencing the fraction of active cells at a given time. (D) Comparison of fraction of failed bursts, where an assembled preinitiation complex produced zero mRNAs before disassembly, as a function of *CTD*_*L*_ by model. Error bars indicate 99% bootstrapped confidence interval. Mean inter-burst duration (E) and burst size (F) as a function of *CTD*_*L*_ and individually varying parameter values, while the others are held constant, as indicated in the first left column of each heatmap. Colormap is artificially fixed to the range [0-20] for visualization purposes and actual numbers are shown in each cell. Model is indicated in the top left corner.

